# Expansin-controlled cell wall stiffness regulates root growth in *Arabidopsis*

**DOI:** 10.1101/2020.06.25.170969

**Authors:** Marketa Samalova, Kareem Elsayad, Alesia Melnikava, Alexis Peaucelle, Evelina Gahurova, Jaromir Gumulec, Ioannis Spyroglou, Elena V. Zemlyanskaya, Elena V. Ubogoeva, Jan Hejatko

## Abstract

Expansins facilitate cell expansion via mediating pH-dependent cell wall (CW) loosening. However, the role of expansins in the control of biomechanical CW properties in the tissue and organ context remains elusive. We determined hormonal responsiveness and specificity of expression and localization of expansins predicted to be direct targets of cytokinin signalling. We found EXPA1 homogenously distributed throughout the CW of columella/ lateral root cap, while EXPA10 and EXPA14 localized predominantly at the three-cell boundaries of epidermis/cortex in various root zones. Cell type-specific localization of EXPA15 overlaps with higher CW stiffness measured via Brillouin light scattering microscopy. As indicated by both Brillouin frequency shift and AFM-measured Young’s modulus, *EXPA1* overexpression upregulated CW stiffness, associated with shortening of the root apical meristem and root growth arrest. We propose that root growth in *Arabidopsis* requires delicate orchestration of biomechanical CW properties via tight regulation of various expansins’ localization to specific cell types and extracellular domains.

## INTRODUCTION

The cell wall (CW) is a fundamental component of plant cells that shapes the plant body, plays key roles in development and growth of organs, movement of solutes and nutrients and protects plants from the environment. CW developmental importance is also recognised in the control of cell differentiation and intercellular communication (Wolf *et al*., 2012). CWs provide the strength, yet they have the ability to expand. Recent studies of growth regulatory networks suggest that the turgor-driven cell expansion is the result of a fine-tuned balance between wall relaxation and stiffening linked by a mechanosensing feedback loop (Braybrook and Jönsson, 2016; Sassi and Trass, 2015; Willis *et al*., 2016). These regulatory networks comprise transcription factors and plant hormones and allow tight control over equilibrium between cell division and differentiation, a process fundamental for growth and development of individual organs in any multicellular organism.

The primary CW consists predominantly of the polysaccharides, cellulose, hemicellulose and pectins. Cellulose microfibers provide the main load-bearing characteristics of the CW, however, the presence of hemicellulose and pectins can alter the viscoelastic properties of the wall matrix (Cosgrove, 2018; Wolf *et al*., 2012). Once the final cell size is reached, a secondary CW can be deposited in specific cell types e.g. xylem tracheary elements (for review see Didi *et al*., 2015) to provide additional mechanical strength. Modulation of CW mechanical properties occurs through the control of biochemical composition as well as the degree and nature of linkages between the CW polysaccharides. Interestingly, wall extensibility may be controlled at limited regions, so called ‘biomechanical hotspots’, where cellulose-cellulose contacts are made, potentially mediated by trace amounts of xyloglucan (Cosgrove, 2014; 2018; 2018b). These relatively limited contact points between cellulose microfibrils may be key sites of a complex process allowing targeted wall expansion, the cell wall loosening.

Expansins, originally described as CW loosening agents during “acid growth” (McQueen-Mason *et al*., 1992), become activated during CW acidification triggered by a number of stimuli through the plasma membrane H^+^-ATPase proton pump (Cosgrove, 2005). Mechanistically, expansins neither possess polysaccharide hydrolytic activity nor change composition of the CW; instead they are proposed to disrupt non-covalent bonds between cellulose and components of surrounding CW matrix, thus relaxing wall stresses and allowing turgor-driven expansion (McQueen-Mason *et al*., 1995; Cosgrove, 2005). *Arabidopsis thaliana* has 36 members of the expansin superfamily (Sampedro and Cosgrove, 2005) that promote CW loosening or are related to the growth of specific cells. *EXPA1* (At1g69530) is known for decades from experiments with beads loaded with purified expansin that induced leaf primordia formation on the shoot apical meristem of tomato (Fleming *et al*., 1997). Apart from leaf organogenesis (Reinhardt *et al*., 1998) and vascular tissue differentiation (Cho and Kende, 1998), expansins are involved in root development and growth (Lee and Kim, 2013; Pacifici *et al*., 2018; Ramakrishna *et al*., 2019), root hair initiation (Cho and Cosgrove, 2002) and seed germination (Sanchez-Montesino, *et al*., 2019; Ribas *et al*., 2019). Interestingly, NbEXPA1 was shown to be plasmodesmata-specific and functions as a novel host factor for potyviral infection (Park *et al*., 2017).

The biomechanical interactions of cells with extracellular matrix have been demonstrated as an important regulator of cell fate specification in animal models (Engler *et al*., 2006). In plants, changes in the CW mechanics are a driving force of growth and development as predicted by a number of biomechanical models (Braybrook and Jonsson, 2016; Geitmann and Ortega, 2009; Haas *et al*., 2020; Hamant and Traas, 2010; Sassi and Traas, 2015). To name a few, *in vivo* chemical modification (demethylesterification) of homogalacturonan by pectin methyl-esterases was shown to be sufficient for the initiation of novel flower and floral organ primordia in *Arabidopsis. Vice versa*, inhibition of homogalacturonan demethylesterification resulted in the inhibition of normal organ formation (Peaucelle *et al*., 2008). Importantly, demethylesterification of homogalacturonan was shown to be associated with an increase in CW plasticity as measured via atomic force microscopy (AFM), suggesting that higher elasticity of CWs might be instructive for newly formed organ primordia (Peaucelle *et al*., 2011). In plants, the importance of biomechanical CW properties has been described mostly in the shoot tissues (Reinhardt *et al*., 1998; Pien *et al*., 2001; Hamant *et al*., 2008; Sampathkumar *et al*., 2014; Landrein *et al*., 2015; Gruel *et al*., 2016; Hervieux *et al*., 2017; Majda *et al*., 2017; Takatani *et al*., 2020). However, the biomechanical interactions associated mostly with the control of CW properties are emerging as an important mechanism guiding also root growth and development (Vermeer *et al*., 2014; Barbez *et al*., 2017; Pacifici *et al*., 2018; Ramakrishna *et al*., 2019; Hurny *et al*., 2020).

Phytohormones including auxins and cytokinins, are key players in growth regulation responses and are thus determinants of plant architecture and CW development. Well known is the role of auxins in the “acid growth theory” (Cleland, 1971; Hager *et al*., 1971), in which auxin induced extrusion of protons into the apoplast activates expansins, leading to CW loosening and growth. Cytokinins were described as factors controlling the equilibrium between cell division and differentiation (Dello Ioio *et al*., 2007, 2008) in the root apical meristem (RAM) by positioning the auxin minimum that triggers the developmental switch (Di Mambro *et al*., 2017). Recently, Pacifici *et al*. (2018) proposed EXPA1 as a direct target of multistep phosphorelay signalling in the cytokinin-regulated cell differentiation in the RAM.

Brillouin light scattering (BLS) is the inelastic scattering of light from inherent or stimulated high frequency acoustic vibrations in a sample, the speed of which is directly related to the elastic modulus of the material (Berne and Pecora, 2000). BLS microscopy is an all optical label-free spectroscopic technique that allows one to spatially map the so-called inelastic Brillouin frequency shift (BFS, ν_B_) with near diffraction limited lateral resolution (Elsayad *et al*., 2019; Prevedel *et al*., 2019) of live cells (Scarcelli *et al*., 2015) and tissue (Elsayad *et al*., 2016). Advances in Brillouin spectrometer design over the last decade (Scarcelli and Yun, 2007) have allowed for studies of live cell and tissue biomechanics at near physiological conditions. In a typical confocal Brillouin microscope, the detector is replaced by a virtually imaged phased-array (VIPA)-based Brillouin spectrometer which acquires an image of the spectrum on the electron multiplying (EM)-CCD camera. The distance of the Brillouin peaks (in GHz) from the central laser frequency is a measure of the local mechanical properties at the confocal volume. Despite probing a distinct elastic modulus in a different frequency regime, the measured BLS has been empirically found to (semi-logarithmically) scale with the lower frequency stiffness defined via the Young’s modulus. BFS can be expected to be higher for “stiffer” samples and smaller for “softer” cells and tissue (Andriotis *et al*., 2019; Gouveia *et al*., 2019; Scarcelli *et al*., 2015). The Young’s modulus is typically measured by atomic force microscopy (AFM). This method is based on micrometer or nanometer tissue compressions (indentations) and was developed to measure precisely the mechanical properties of CWs in developing organs and across entire tissue regions (Peaucelle, 2014). The measured stiffness (resistance to deformation/ indentation) is defined by the measurement of an indentation modulus that best describes the elasticity of the scaffolding of the CW of the tissue. AFM can be also used to image CW surface topology at high resolution to detect individual cellulose microfibrils (app. 3 nm in diameter, Zhang *et al*., 2014) and can be carried out under water, allowing imaging of CW in a near-native state.

In this work, we set out to localise EXPA1 and its homologues EXPA10, EXPA14 and EXPA15 and describe the relationship between expression of expansins and the mechanical properties of the CW during root cell differentiation. To quantitate the changes in CW biomechanics, we introduce mechano-optical contrast (MOC) as a new measure of mechanical properties for various biological structures measured via Brillouin light scattering and confirm the results using more established atomic force microscopy (AFM).

## RESULTS

### Cytokinins and auxins control *EXPAs* transcription in the *Arabidopsis* root

For our study we selected expansins AtEXPA1 (EXPA1, *At1g69530*), AtEXPA10 (EXPA10, *At1g26770*), AtEXPA14 (EXPA14, *At1g69530*) and AtEXPA15 (EXPA15, *At2g03090*), previously suggested to be under hormonal control (Lee *et al*., 2007; Bhargava *et al*., 2013; Pacifici, *et al*., 2018; Ramakrishna *et al*., 2019).

According to published data (Zubo *et al*., 2017; Pacifici *et al*., 2018; Taniguchi *et al*., 2007; Xie *et al*., 2018), *EXPA1* was supposed to be the direct target of cytokinin-responsive ARABIDOPSIS RESPONSE REGULATOR 1 (ARR1) and its homologues ARR10 and ARR12 (Figure 1A). EXPA1 responsiveness to auxin could be potentially regulated by AUXIN RESPONSE FACTOR 5 (ARF5) since the corresponding DNA affinity purification (DAP)-sequencing peaks (O’Malley *et al*., 2016) are located in its promoter (Figure 1B). To confirm the hormonal regulations over *EXPA1*, we quantified *EXPA1* transcripts using reverse transcription quantitative real-time PCR (RT qPCR) in wild-type (WT) *Arabidopsis* seedlings treated with 5 μM 6-Benzylaminopurine (BAP) and 5 μM 1-naphthaleneacetic acid (NAA, Figure 1C). With the cytokinin treatment, transcript levels were transiently and rather moderately (3-4 times compared to the mock-treated control) upregulated during the 4-hour time span tested; similar results were obtained using *trans*-zeatin (tZ, data not shown). Compared to that, *EXPA1* responded more distinctly to the auxin treatment and its transcript level was increased continuously up to 5–10 fold at 4h. However, no increase in *EXPA1* promoter activity was detected in a pEXPA1::nls:3xGFP transcriptional fusion line (Ramakrishna *et al*., 2019) after both cytokinin and auxin treatment (Figure 1-figure supplement 1) neither in RAM nor TZ.

**Figure 1:**
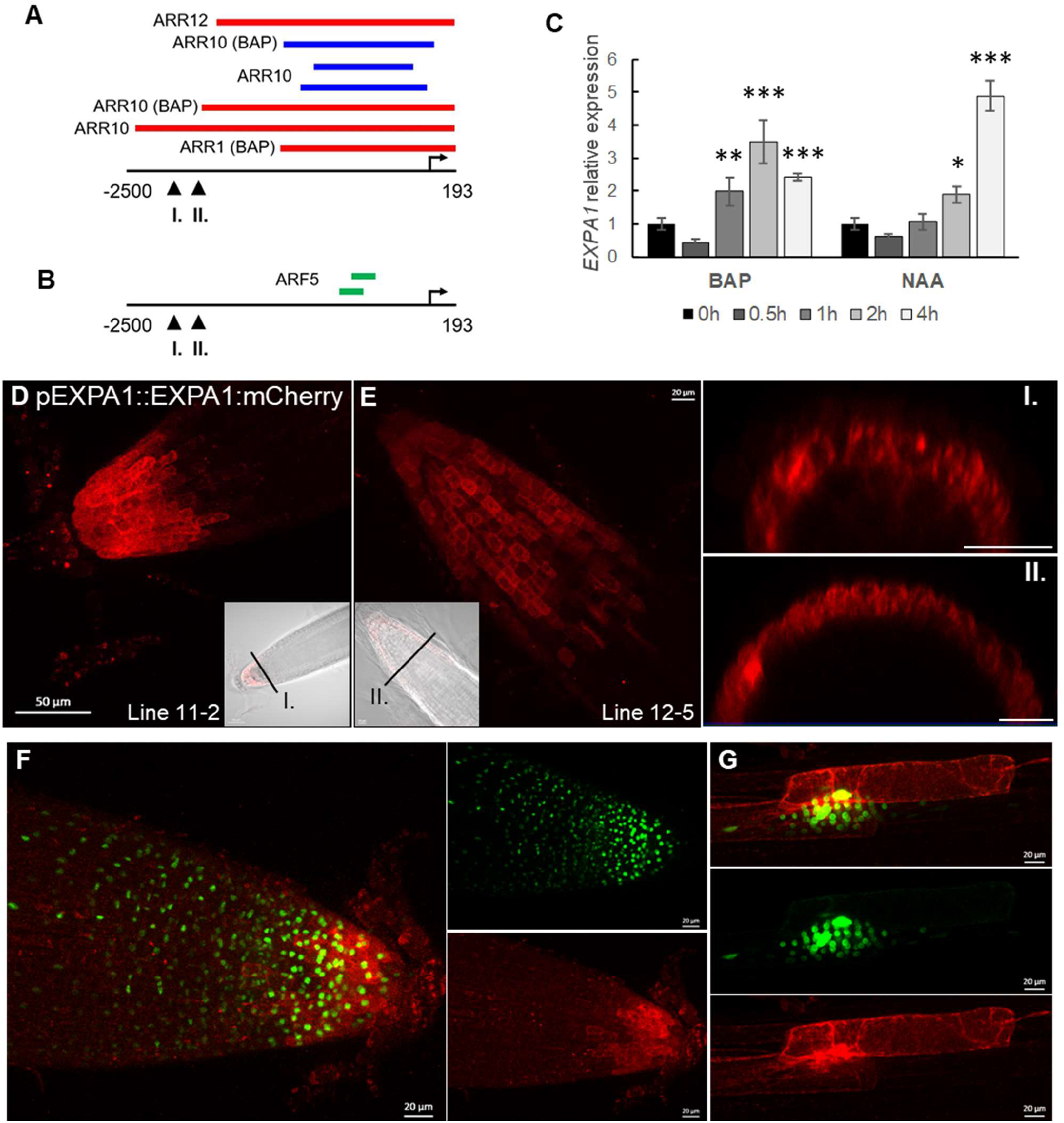
Transcript profiling of *EXPA1* in response to hormones, *EXPA1* promoter activity and EXPA1 localization. **(A)** *EXPA1* promoter analysis identifies ChIP-seq- and DAP-seq-derived binding events for transcription factors involved in cytokinin and **(B)** auxin signalling pathways. Red, blue and green colours depict the peaks from Xie *et al*., 2018, Zubo *et al*., 2017 and O’Malley *et al*., 2016, respectively. The coordinates are represented relative to the transcription start site marked by the arrow. The arrowheads indicate the 5’-end of the pEXPA1 promoter in (I.) this publication and (II). Pacifici *et al*., 2018. **(C)** Quantitative real-time PCR of roots of 7-day old *Arabidopsis* WT seedlings treated with 5 μM BAP or 5 μM NAA for 0.5h, 1h, 2h and 4h. The transcript abundance of EXPA1 is double normalized to UBQ10 and mock-treated controls. The experiment was repeated twice with 3 replicas of each sample, error bars represent SD. Statistically significant differences at alpha 0.05 (*), 0.01 (**) and 0.001 (***) are shown. **(D, E)** Z-stack projections of pEXPA1::EXPA1:mCherry fusion localised in the LRC and columella of two independent singlecopy transgenic lines 11-2 (D) and 12-5 (E) and their transversal xz optical sections (I. and II.) as indicated by the black lines in the transmitted-light micrograph inserts shown as a single optical section. **(F, G)** Z-stack projections of F1 line pEXPA1::EXPA1:mCherry (11-2) crossed with pEXPA1::nls:3xGFP illustrating a similar pattern of EXPA1 expression by GFP (green) and mCherry (red) signals in RAM (F) and lateral root primordium (G). Scale bars correspond to 20 μm except in D, where it corresponds to 50 μm.

Based on our *in silico* analysis, *EXPA10, EXPA14* and *EXPA15* (Figure 2-figure supplement 1) might also be direct targets of cytokinin-activated type-B ARRs. In line with that, both *EXPA14* and *EXPA15* were upregulated by cytokinins; nonetheless, in contrast to previous report (Pacifici *et al*., 2018), no positive response has been detected in case of *EXPA10* (both BAP and tZ, Figure 2 and data not shown).

**Figure 2:**
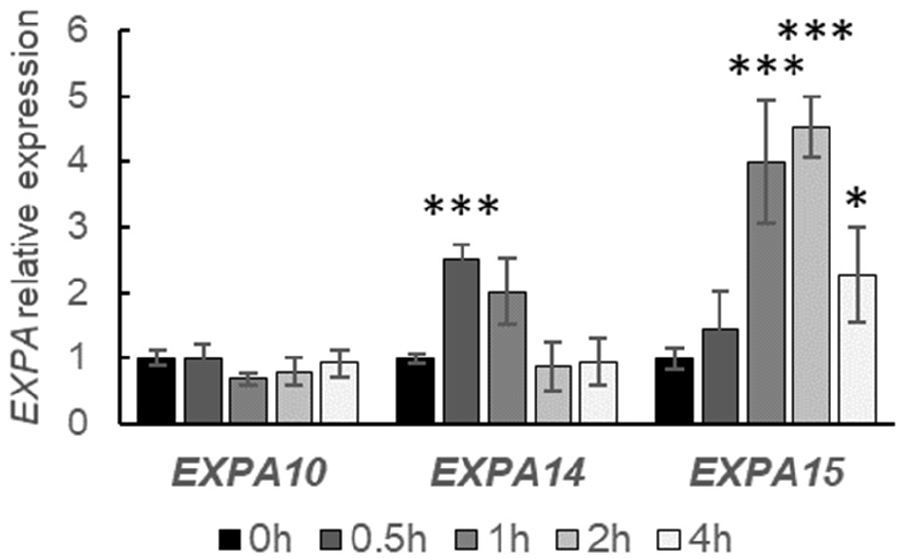
Transcript profiling of *EXPA10, EXPA14* and *EXPA15* in a response to cytokinin treatment. Quantitative real-time PCR of roots of 7-day old *Arabidopsis* WT seedlings treated with 5 μM BAP for 0.5h, 1h, 2h and 4h. The transcript abundance of the *EXPAs* is double normalized to UBQ10 and mock-treated controls. The experiment was repeated twice with 3 replicas of each sample, error bars represent SD. Statistically significant differences at alpha 0.05 (*) and 0.001 (***) are shown.

Altogether, our data suggest moderate and transient *EXPA1* and *EXPA14* upregulation by cytokinins, stronger response was seen in case of auxin- and cytokinin-mediated upregulation of *EXPA1* and *EXPA15*, respectively. However, no clear effect of exogenous hormone application was detectable for *EXPA10*.

### Expansins localise to the root CW in a specific pattern

Previously, expansins were shown to be localised in the CW by immunogold labelling of CWs and Golgi-derived vesicles using antibody against α-expansin (Cosgrove *et al*., 2002). However, so far attempts to visualise expansins in the CW of living plants by a translational fusion with a green fluorescent protein (GFP) failed (Pacifici *et al*., 2018), perhaps due to high sensitivity of GFP to the low pH environment. Therefore, we created translational fusions of EXPA1, EXPA10, EXPA14 and EXPA15 with a red fluorescent protein mCherry (Shaner *et al*., 2004) under the control of native promoters and confirmed their CW localisation in a highly tissue-specific manner in roots (Figures 1 and 3).

**Figure 3:**
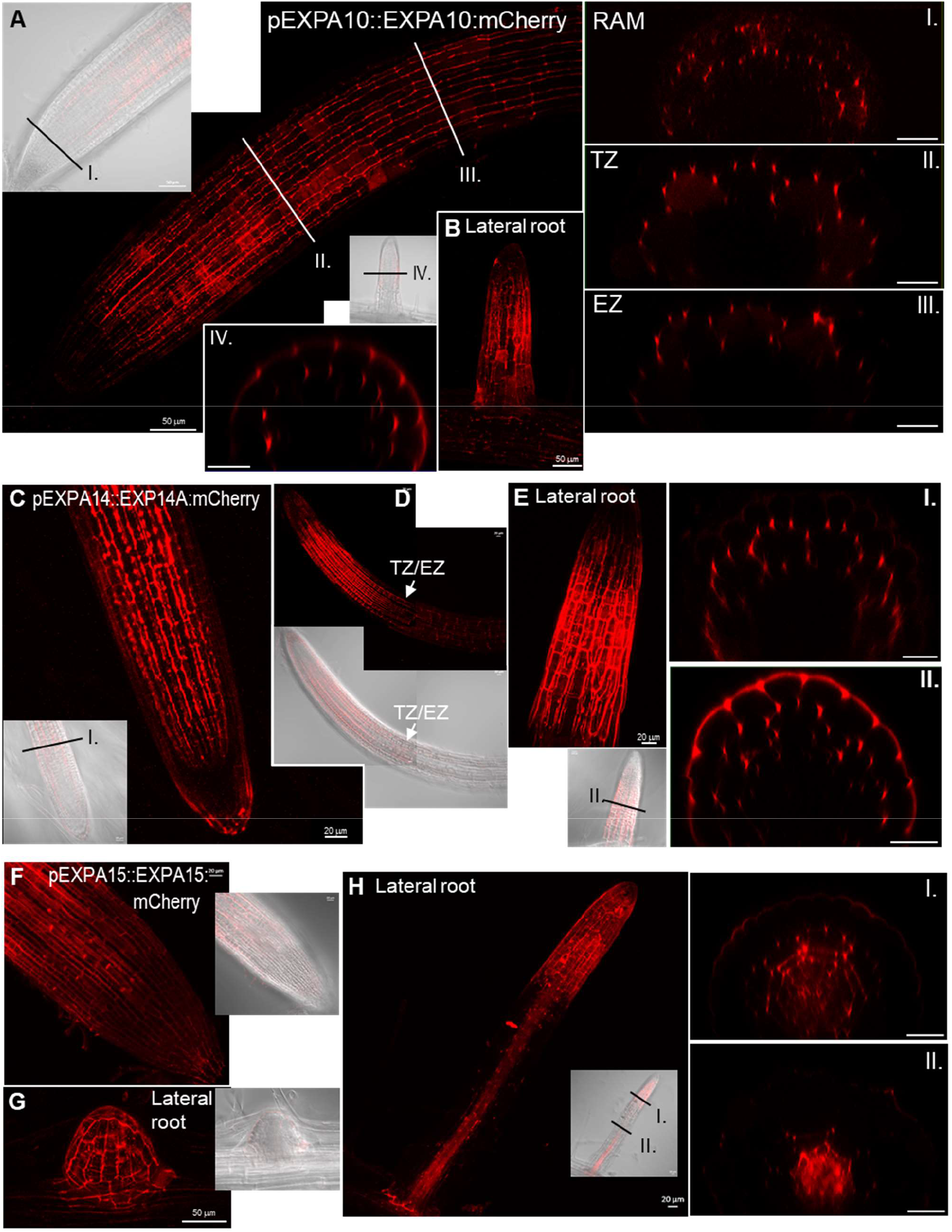
EXPA10, EXPA14 and EXPA15 reveal specific localization in the root tip. **(A, B)** pEXPA10::EXPA10:mCherry in a primary (A) and lateral (B) root and transversal xz optical sections (I.-IV.) through the RAM, transition (TZ) and elongation (EZ) zones and the lateral root as indicated by the black and white lines in the longitudinal plane views. **(C-E)** pEXPA14::EXPA14:mCherry in a primary (C, D) and lateral (E) root and transversal xz optical sections (I. and II.) as indicated by the black lines in the longitudinal plane views; the white arrows point to the TZ/EZ boundary. **(F)** pEXPA15::EXPA15:mCherry in RAM, **(G)** an emerging lateral root, **(H)** a lateral root and its transversal xz optical sections (I. and II.) as indicated by the black lines. Z-stack projections from optical sections taken from the top of the root to the longitudinal plane are shown in all longitudinal plane views. Transmitted-light micrograph inserts show a single optical section. Scale bars correspond to 20 μm except in A, B, G, where they correspond to 50 μm.

In *Arabidopsis* root, *EXPA1* revealed the strongest expression in the columella and lateral root cap (LRC) of both the main and lateral roots (Figure 1D-G). Interestingly, the cells immediately surrounding developing lateral roots and primordia were also strongly expressing *EXPA1* (Figure 1G). These results were confirmed using an independent transcriptional *pEXPA1::nls:3xGFP* fusion line (Ramakrishna *et al*., 2019) crossed into the mCherry line background (Figure 1F and 1G). Occasionally, very weak *EXPA1* promoter activity was detectable in the root transition zone (TZ)/elongation zone (EZ) boundary (Figure 1 – figure supplement 2A); however, no EXPA1:mCherry was detectable there (Figure 1 – figure supplement 2B).

*EXPA10* was visibly expressed in the cortex layer of the primary root from the meristematic zone up (proximally) to the first lateral roots (Figure 3A). Unlike *EXPA1, EXPA10* is not expressed in the LRC. Interestingly, in contrast to a rather homogenous distribution of EXPA1 throughout the CW surrounding the LRC/columella cells, we observed distinct “spotty” localisation of EXPA10 dominantly in the cortex/endodermis and cortex/epidermis three-cell boundaries that was particularly visible on the cross-sections of the roots (Figure 3A, insets I.-III.). This kind of unequal distribution of EXPA10 in the three-cell boundaries was occasionally detectable also in the longitudinal plane view of the root cortex cell files. Here, the positions of EXPA10 localization maxima do not seem to overlap with cellulose deposition as detected using calcofluor white staining of fixed roots (Figure 3 – figure supplement 1, white arrows). In the lateral roots, the EXPA10:mCherry fusion localised predominantly in the epidermis and cortex layers of the transition/elongation zones (Figure 3B and inset IV.).

EXPA14:mCherry fusion localises in the cortex layer of the RAM up to the TZ/EZ boundary, proximally from which the signal disappears (Figures 3C, D). The distinct pattern of strong accumulation of the protein in the apoplastic space at the boundary of three cells (insets for Figure 3C and 3E, Figure 3-figure supplement 2A) resembles the one observed for EXPA10:mCherry fusion. In the lateral roots, *EXPA14* is also strongly expressed in the transition/elongation zones (Figure 3E). However, in contrast to the situation in the main root, in the lateral root EXPA14 locates not only to cortex, but also to the epidermal cell layers (Figure 3E inset II.).

EXPA15:mCherry is localised in the epidermis of RAM (Figure 3F) and emerging lateral roots (Figure 3G, Figure 3 – figure supplement 2B) in a relatively uniform pattern; however, the “spotty” pattern at three-cell boundary was apparent in the more internal cortex/endodermis. Proximally from the meristematic zone, EXPA15 re-localises into deeper (vasculature) layers, again revealing rather homogenous distribution (Figure 3H insets I.-II.).

Worth of note, in contrast to homogenously distributed EXPA1 and (partially) EXPA15, EXPA10 and EXPA14 seem to be localized dominantly in the longer (parallel with the longitudinal root axis) walls of elongated cells, particularly in the RAM of the main root (Figure 3 – figure supplement 3).

To confirm the extracellular localisation, we activated the (naturally very weak) expression of *EXPA1:Cherry* in all plant tissues (Figure 4) using the dexamethasone (Dex) inducible system pOp6/LhGR (Craft *et al*., 2005; Samalova *et al*., 2005; Samalova *et al*., 2019). After both long (7 days) and short (24 h) Dex induction, the fusion protein accumulated in the cell periphery/apoplastic space in roots but was also visible in the transit through the secretory pathway from the endoplasmic reticulum (ER) to the CW. However, since the resolution of a confocal microscope does not allow to distinguish between CW and plasma membrane localisation, we treated the roots with 10% sorbitol to allow for plasmolysis. Figure 4G shows that unlike the plasma membrane marker (Figure 4-figure supplement 1), EXPA1:mCherry remained located at the outer edges of the cells, suggesting that EXPA1 is indeed localised in the cell wall. Importantly, the CW localization pattern we observed in case of the Dex-induced pRPS5A>GR>EXPA1:mCherry line was still resembling the homogenous distribution we observed in case of EXPA1:mCherry driven by its natural promoter in the lateral root cap tissue (compare Figures 1 and 4). This is suggesting that the CW localization pattern of EXPA1 is independent of the cell type and the level of expression.

**Figure 4:**
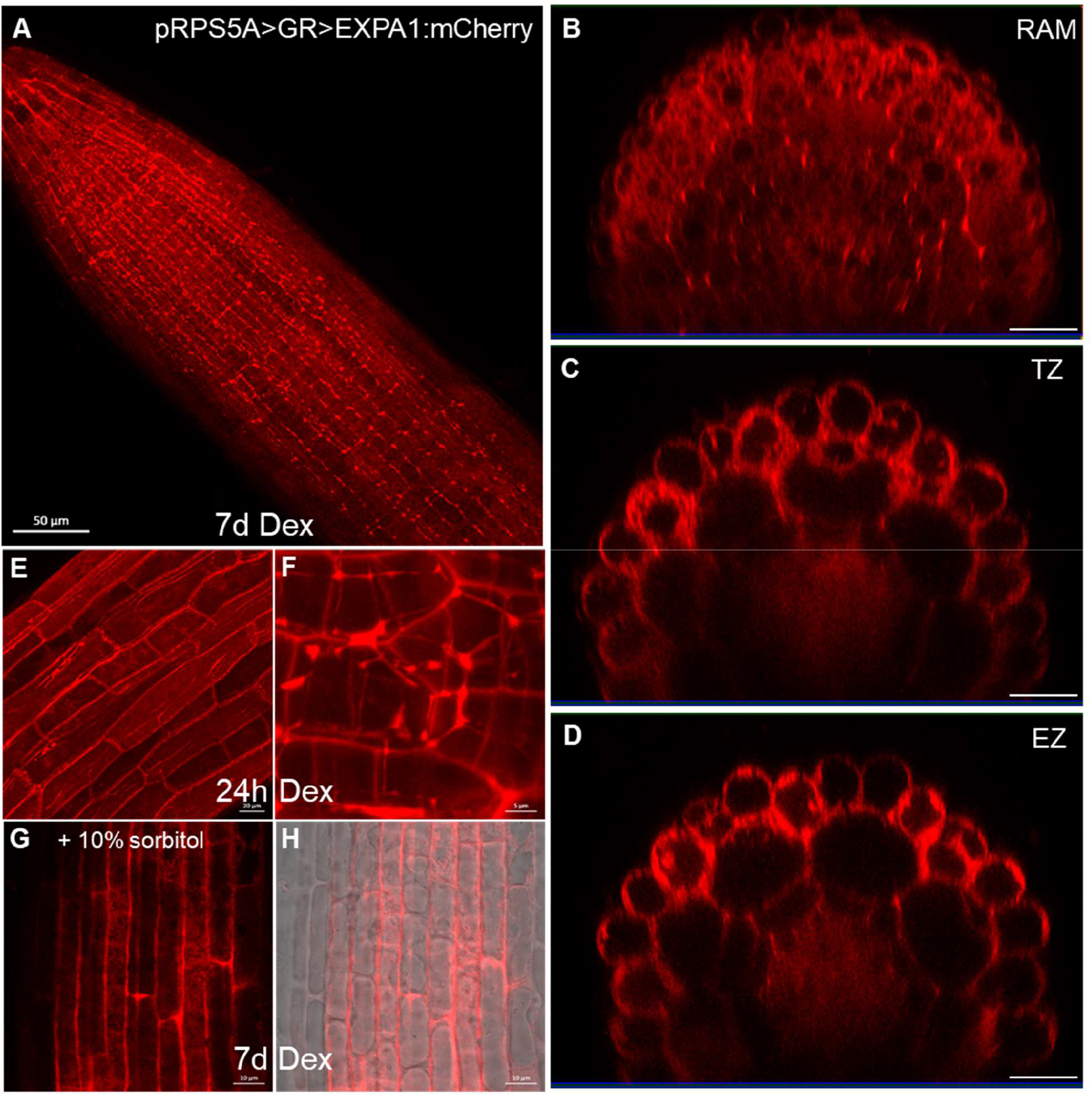
Overexpressed EXPA1 localizes to the cell wall. Z-stack projections of EXPA1:mCherry fluorescence in pRPS5A>GR>EXPA1:mCherry seedlings induced by Dex on a solid MS medium for 7d in a primary root **(A)** and its xz optical cross-sections through RAM **(B)**, TZ **(C)** and EZ **(D)**. **(E, F)** pRPS5A>GR>EXPA1:mCherry seedlings induced by Dex in a liquid MS medium for 24h and imaged EZ of a primary root (E) and a lateral root (F). **(G,H)** pRPS5A>GR>EXPA1:mCherry seedlings induced as in (A) and after 10 min-treatment of the primary root with 10% sorbitol. Fluorescence channel (G) and its overlay with transmitted light (H). Scale bars correspond to 20 μm except in A is 50 μm, in F is 5 μm and in G and H are 10 μm.

To conclude, all assayed expansins show distinct expression and localization patterns. The differences in the localization pattern between EXPA1 revealing homogenous distribution all around the cell and the “spotty” localization of EXPA10, EXPA14 and (partially) EXPA15 implies differential roles of individual EXPAs in the control over root CW properties.

### Introducing mechano-optical contrast for Brillouin-based imaging of biological samples

To investigate the biomechanical properties on the sub-cellular level we used Brillouin light scattering (BLS) microscopy. The Brillouin frequency shift (ν_B_) is proportional to the acoustic phonon velocity, which is in turn proportional to the square root of the high frequency longitudinal elastic modulus (M). As such, the Brillouin frequency can serve as a proxy for the mechanical properties of the sample. In particular, M is closely related to the compressibility of the sample, and has empirically been observed to scale semi-logarithmically with the Young’s modulus (E) as measured by AFM in diverse samples including live cells (Scarcelli *et al*., 2015). An exact calculation of M requires knowledge of the ratio *n*^2^/ρ where *n* and ρ are the refractive index and mass density in the probed region of the sample respectively. While it can by virtue of the Lorentz-Lorenz (LL) relation often be assumed that *n*^2^ will scale with ρ (Zhao *et al*., 2011) such that explicit knowledge of the ratio *n*^2^/ρ at each probed region is not required, the validity of the LL-relation in complex multicomponent structures including the CW cannot be rigorously justified. As such, we present results in terms of a dimensionless frequency shift we term the *Mechano-Optical Contrast* (MOC), 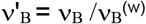, where **ν**_B_^(w)^ is the measured BLS frequency of distilled water (Antonacci *et al*., 2020). As is the case for the Brillouin frequency shift, the MOC will scale as the square root of M, with the advantage that it is independent of the probing wavelength, can correct for slight variations in temperature between measurements and allows for the more straight forward comparisons of measurements between instruments employing different probing wavelengths.

### *EXPA1* overexpression stiffens root cell wall

To characterize its importance in the control of biomechanical CW properties, we overexpressed *EXPA1* (without any tag) and generated pRPS5A>GR>EXPA1 Dex-inducible lines (8-4 and 5-4) using the pOp6/LhGR system as above. Representative 2D Brillouin frequency shift maps (Figure 5A) display the BLS shift in the CW of plants overexpressing *EXPA1* before and after induction. We quantified the MOC in roots (early EZ) of 7-day old *Arabidopsis* WT and *EXPA1* overexpressing seedlings (line 8-4) grown on MS media pH 5.8 or pH 4 (Figure 5B and C, respectively) with or without Dex induction. From the technical reasons (to obtain sufficient overlap of point spread function with cell wall and hence good CW signal) and the expression profile of assayed *EXPA* genes (epidermis and/or cortex), we focused on the longitudinal inner epidermal CWs (epidermis/cortex boundary). The plants overexpressing *EXPA1* showed a higher MOC (longitudinal elastic modulus, *vide supra*) for the root CWs on both pH media, suggesting that their CWs are stiffer. The pRPS5A>GR>EXPA1:mCherry lines induced on Dex also displayed higher MOC, however not significantly different from the non-induced plants (Figure 5 - figure supplement 1A), perhaps due to lower expression levels of the *EXPA1* (Figure 5 - figure supplement 2). Importantly, we have detected increase in the MOC/CW stiffness even in response to short-term Dex-mediated *EXPA1* upregulation. For the strong expression line pRPS5A>GR>EXPA1 (8-4), the statistically significant change was observed as soon as 3 h after *EXPA1* induction (Figure 5D, E) on media at both pH tested (4 and 5.8). In case of a weaker expression line pRPS5A>GR>EXPA1:mCherry (1-3), the significant increase in MOC was detected later, after 6 h of induction and only on the media with pH 4.0 (Figure 5 - figure supplement 1B). To exclude the unspecific/side effects of gene overexpression, we determined the spatial map of cell stiffness using fluorescence emission–Brillouin imaging (Elsayad *et al*., 2016) in the *Arabidopsis* root at larger area in a pEXPA15::EXPA15:mCherry line, revealing stronger mCherry signal when compared to pEXPA1::EXPA1:mCherry. The regions of higher Brillouin frequency shift correlated well with *EXPA15* expression domain (Figure 6), suggesting the role of EXPA15 in the control of CW stiffness. On the other hand, we did not detect any changes in the CW stiffness in the root TZ of *expa1-1* knock-out line (Pacifici *et al*., 2018) when compared to WT (Figure 5 - figure supplement 3).

**Figure 5:**
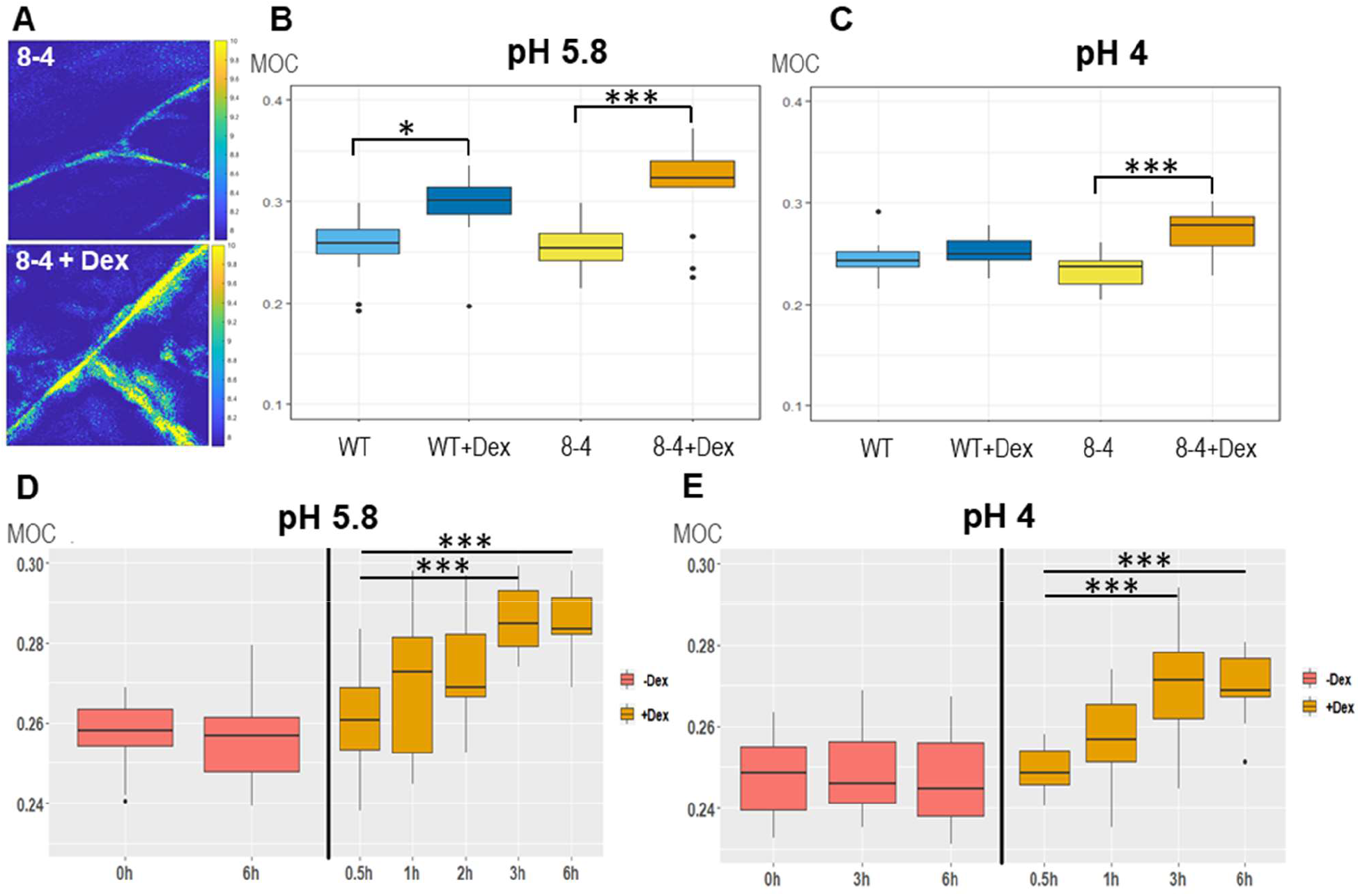
EXPA1 overexpression stiffens cell walls when measured via Brillouin frequency shift. **(A)** Representative images of 2D (*xy*) Brillouin frequency shift (BFS) maps in root cells of 7-day old *Arabidopsis EXPA1* overexpressing seedlings of pRPS5A>GR>EXPA1 (line 8-4) grown on MS media pH 5.8 +/- Dex. BFS expressed as Mechano-Optical Contrast (MOC) was determined in roots of WT and the 8-4 line grown on MS media **(B)** +/- Dex pH 5.8, **(C)** +/- Dex pH 4, **(D)** induced in liquid MS media pH 5.8 for 0.5h – 6h, **(E)** induced in liquid MS media pH 4 for 0.5h – 6h; DMSO was used in -Dex treatments. Medians shown are from at least 4 seedlings and 10 measurements in each category. Statistically significant differences at alpha 0.05 (*) and 0.001 (***) are shown.

**Figure 6:**
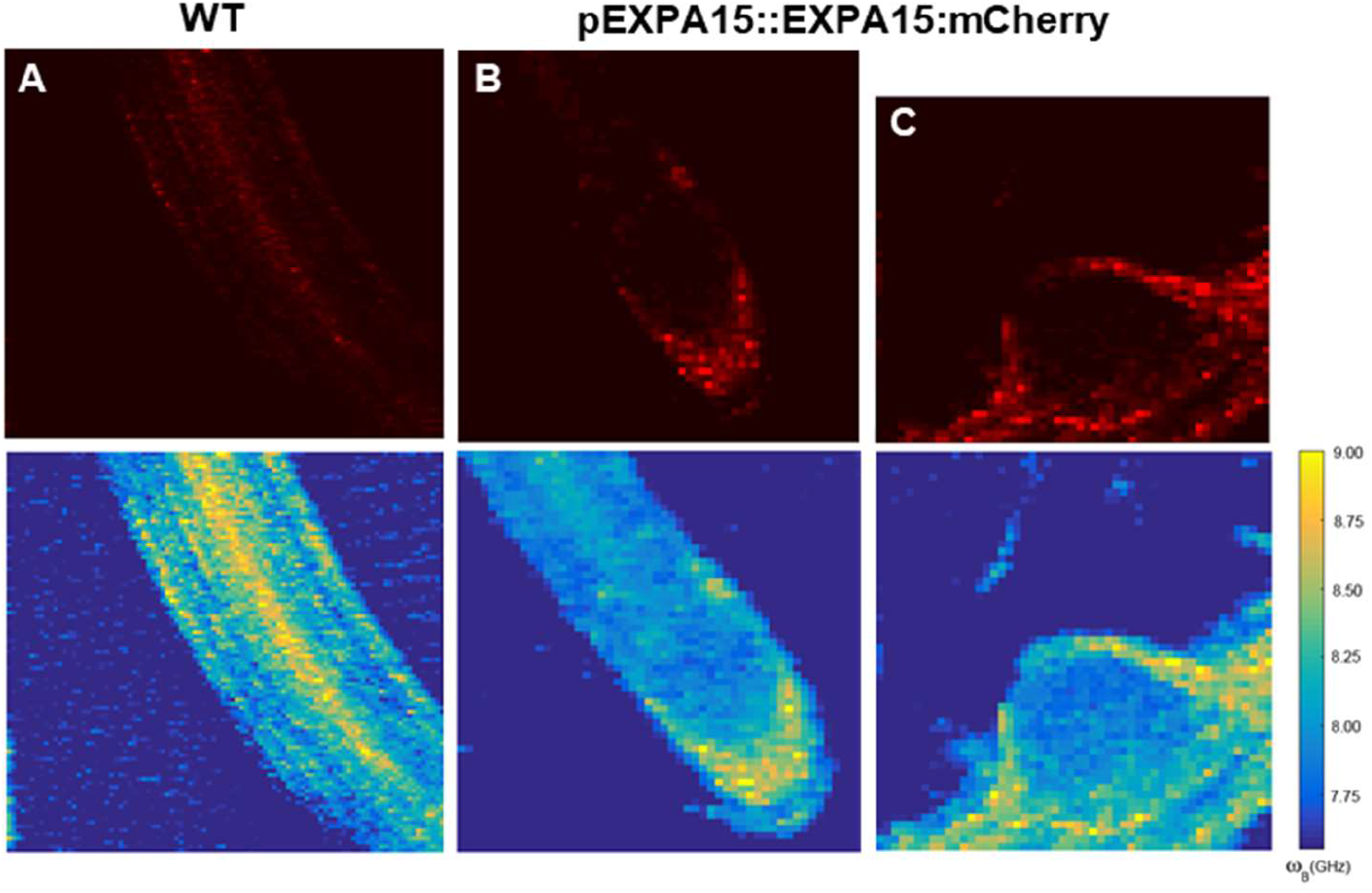
EXPA15 localization overlaps with higher cell wall stiffness in the root tip as determined using fluorescence emission-Brillouin scattering. Fluorescence images (top raw, in red) and Brillouin frequency shifts (BFS, bottom raw, false color-coded) of 7-day old *Arabidopsis* root of **(A)** WT and pEXPA15::EXPA15:mCherry **(B)** a primary root and **(C)** an emerging lateral root. The fluorescent signal corresponds to EXPA15 expression and overlaps with higher BFS.

Since the refractive index (RI) in cells directly correlates with the mass content, we applied quantitative cell tomography (employing a holotomographic microscope) to measure the RI directly in the *Arabidopsis* roots in water. Representative maximum intensity projections of RI tomograms are shown in Figure 7A. The quantitative data analysis confirmed that there are no statistically significant differences across all genotypes and treatment performed in both longitudinal (upper graphs) and transverse (lower graphs) cell walls of the early elongating cells in *Arabidopsis* roots grown at both pH 4 and 5.8 (Figures 7B, C).

**Figure 7:**
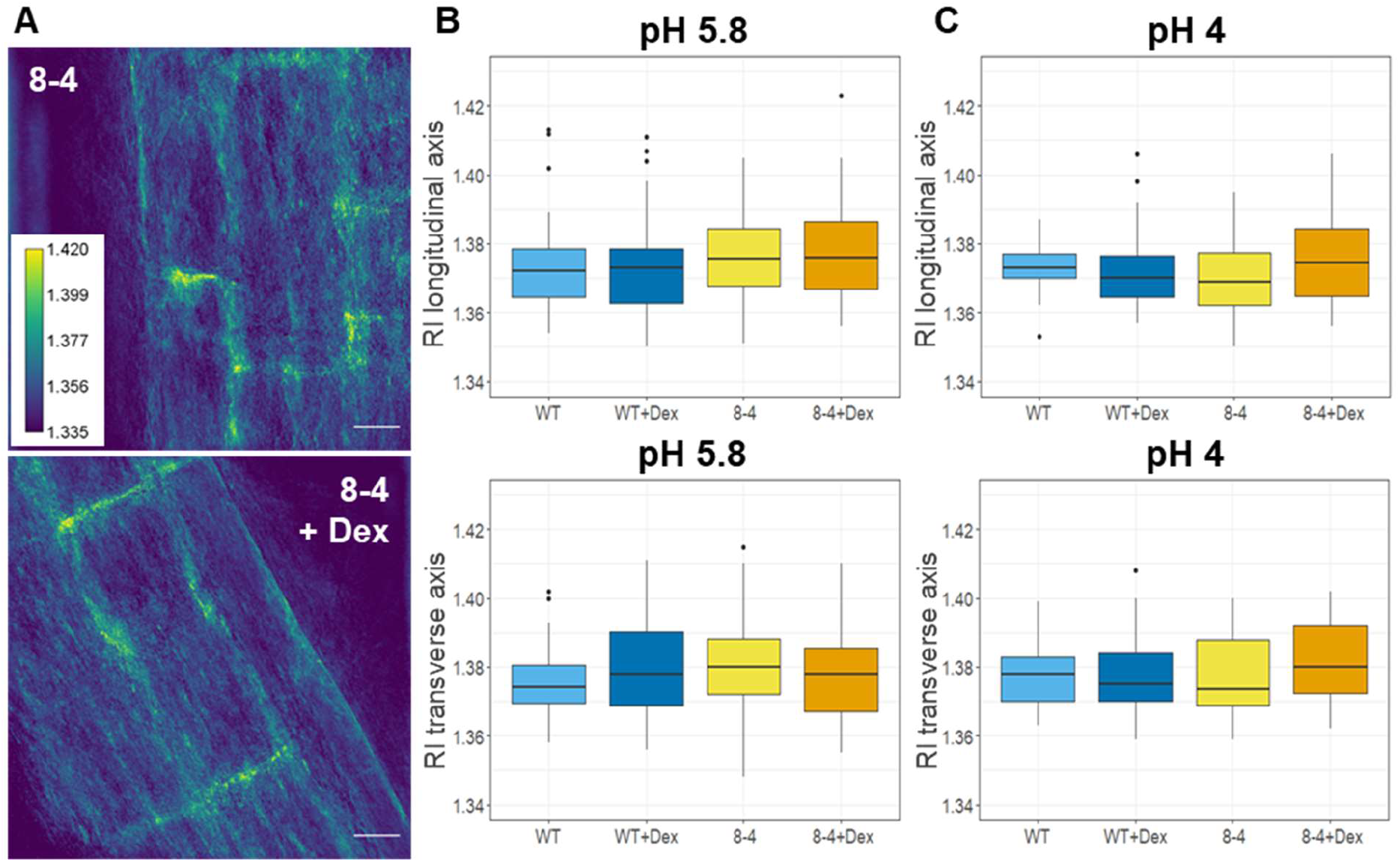
Refractive index measurements of *Arabidopsis* root cell walls reveals comparable cell wall density in wild type and *EXPA1* overexpressing roots. **(A)** Refractive index tomograms (maximal projections) of root cells of 7-day old *Arabidopsis* of *EXPA1* overexpressing seedlings pRPS5A>GR>EXPA1 (line 8-4) grown on MS media pH 5.8 +/- Dex. Scale bars indicate 20 μm. The graphs show RI measurements in water (RI 1.330) of roots of WT and the 8-4 line grown on MS media **(B)** +/- Dex pH 5.8, **(C)** +/- Dex pH 4. Medians from minimum of 6 seedlings and 30 measurements in each category of longitudinal (upper) and transverse (lower) CW axis are shown. There are no statistically significant differences between the genotypes and treatments used.

To directly measure the “stiffness” of root CWs, we used atomic force microscopy (AFM). AFM-based microindentations apply precise known forces on a cell through a cantilever and give a deformation value to extract cell Young’s modulus (Peaucelle, 2014; Peaucelle *et al*., 2015). In a complex structure of plant tissues, at small deformation the force to deform material is proportional to the area of indentation allowing the determination of a coefficient of proportionality that is named “apparent Young’s modulus” (Peaucelle, 2014). This coefficient depends on the speed of deformation and mechanical characteristics of the sample. Representative heat maps of the apparent Young’s modulus (E_A_) show clear differences in E_A_ measured in the CW of root early EZ in 7-day old *Arabidopsis* WT and *EXPA1* overexpressing seedlings (Fig. 8A). The data quantification confirms that the Dex-induced EXPA1 associates with significantly stiffer roots (P<0.001) on growth media at pH 5.8 or pH 4 (Figure 8B and C, respectively). The stiffening effect of *EXPA1* overexpression thus seems to be observable at indentation speed of seconds and at the GHz through the Brillouin technique.

**Figure 8:**
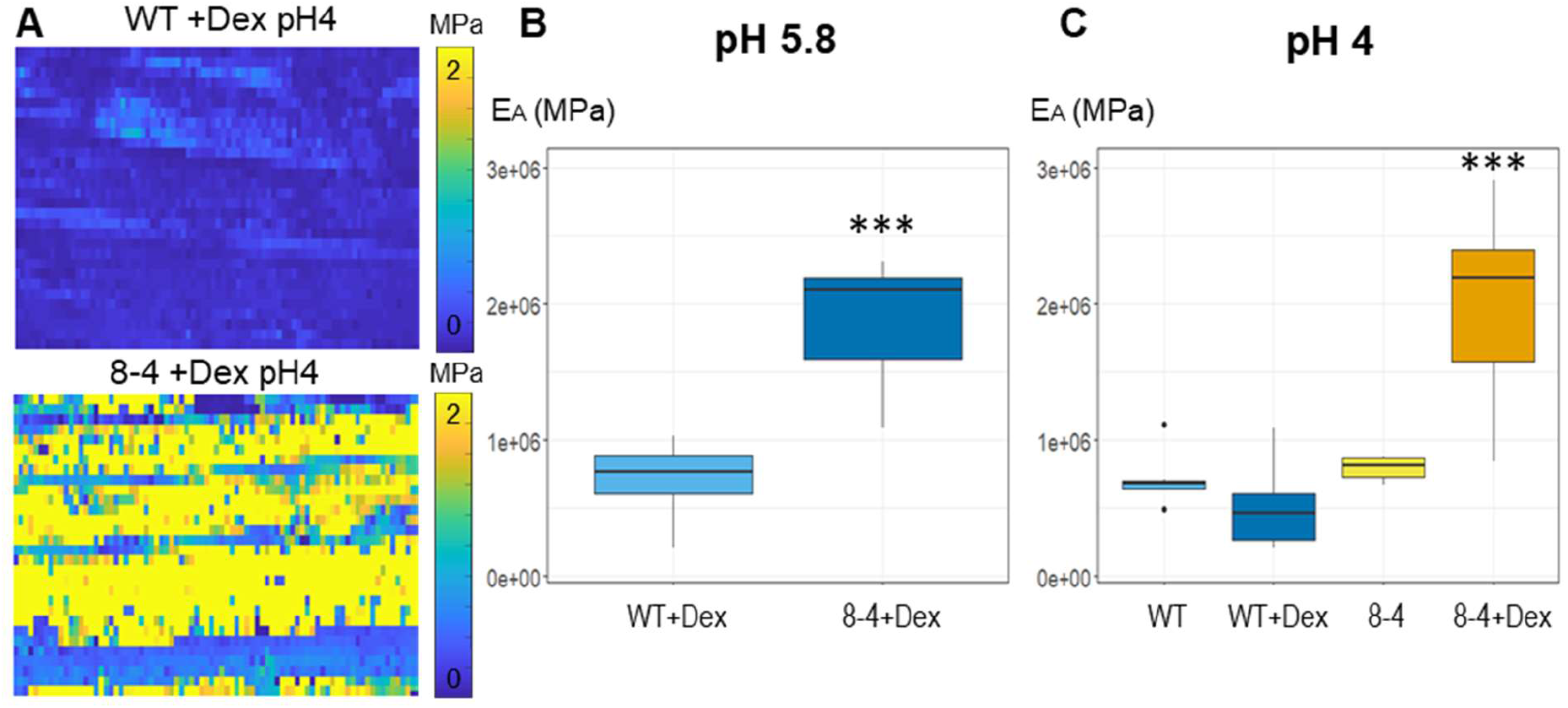
EXPA1 overexpression stiffens cell walls when determined using atomic force microscopy. **(A)** Representative maps of the apparent Young’s modulus (E_A_) of root cells of 7-day old *Arabidopsis* WT and *EXPA1* overexpressing seedlings (line 8-4) grown on MS media pH 4 plus Dex, showing differences in E_A_ (representative of >50). The E_A_ maps are presented as heat maps, with their respective scales, and show data from two successive maps of 60 × 80 and 60 × 80 force scans. Each pixel in the E_A_ map represents the E_A_ calculated from a single force-indentation curve, and each map consists of 4,800 data points. Images are 100 μm in length. Graphs are presenting the E_A_ of the roots as in (A) grown on MS media +/- Dex **(B)** pH 5.8 and **(C)** pH 4. The E_A_ plotted on the graphs was determined by sampling data points within the area of interest. Medians shown are from minimum of 6 measurements in each category. Statistically significant differences at alpha 0.001 (***) are shown.

To wrap up, overexpression of *EXPA1* results into stiffening of the CWs measured at the TZ/EZ boundary using both Brillouin light scattering and AFM. Interestingly, even the natural EXPA15 expression seems to associate with cells revealing higher stiffness within the *Arabidopsis* root tip.

### *EXPA1* overexpression downregulates root growth by reducing RAM size

We examined the phenotype of WT and *EXPA1* overexpressing seedlings (pRPS5A>GR>EXPA1 lines 5-4 and 8-4) grown on Dex continuously for 1 week. The Dex-induced plants had significantly reduced length of roots by 25-30% (Figures 9A, B). The reduction was further enhanced to 40-73% when the pH of the growth media was dropped from 5.8 to 4. A detailed examination of the RAM together with the TZ revealed that the size was significantly reduced by 18% and 29% for the line 8-4 grown on media with normal (5.8) and acidic (4) pH, respectively (Figures 9C). Similarly, the number of the cells (counted in the cortex layer from the quiescent centre to the first elongated cell) was significantly reduced (Figure 9D). However, the ratio size/number of cells remained the same for each line with or without Dex induction (Figure 9E), suggesting that the number of the cells, but not the cell length is reduced in the smaller roots.

**Figure 9:**
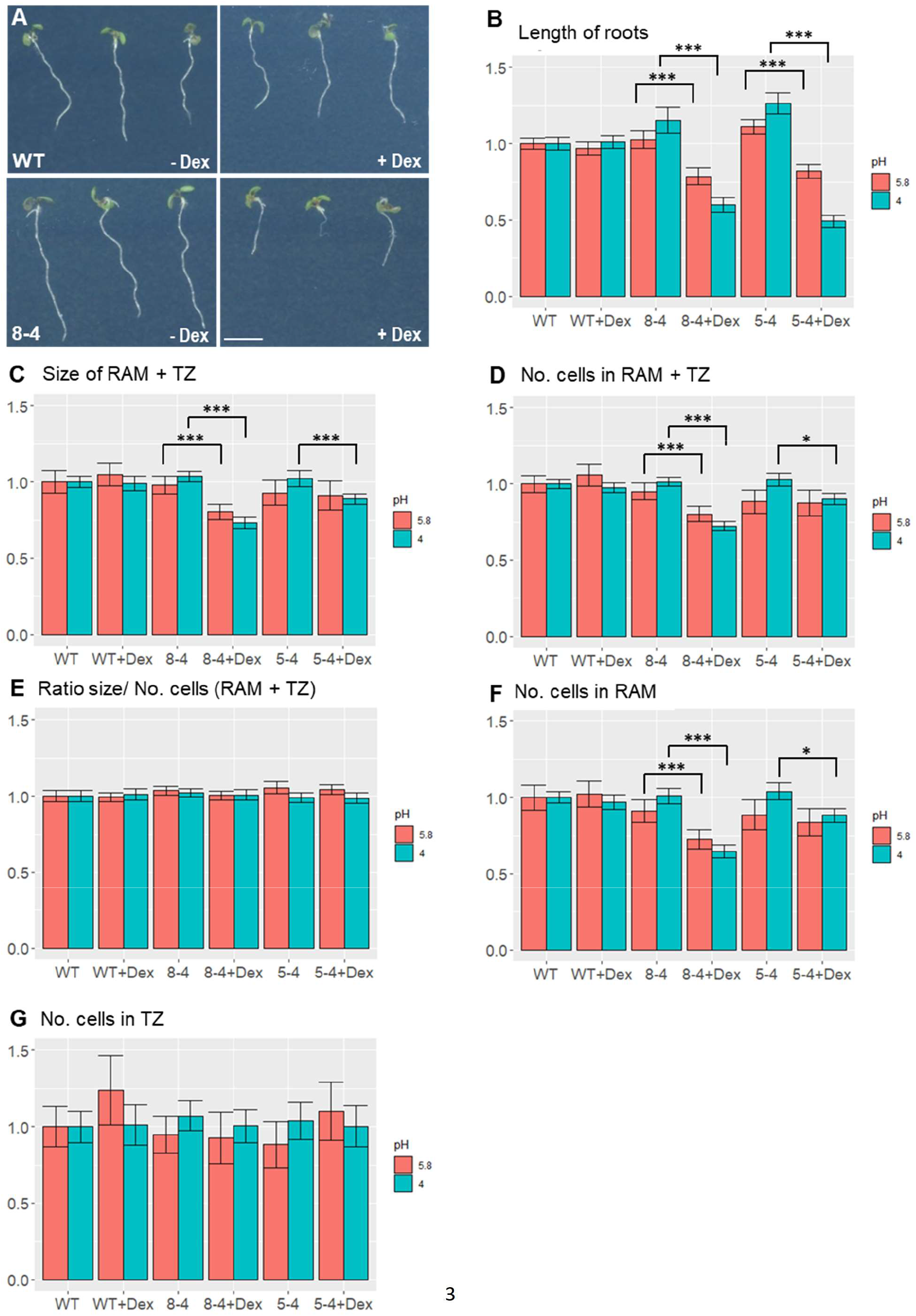
*EXPA1* overexpression reduces root growth via shortening of the root apical meristem. **(A)** 7-day old *Arabidopsis* seedlings of WT (top row) and pRPS5A>GR>EXPA1 line 8-4 (bottom row) grown on MS media pH4 supplemented either with DMSO (-Dex) or dexamethasone (+Dex) The scale bar is 5 mm. **(B)** Length of roots, **(C)** size of RAM + TZ, **(D)** the total number of cells (No. cells) in RAM + TZ, **(E)** the ratio of size/ No. of cells **(F)** No. cells in RAM, **(G)** No. cells in TZ of two independent EXPA1 overexpressing lines 8-4 and 5-4 grown on MS media +/- Dex pH 5.8 and pH 4 relative to WT. Each experiment was repeated at least three times with minimum of 10 seedlings in each category, error bars represent 95% confidence interval. Statistically significant differences within genotypes and treatments at alpha 0.05 (*) and 0.001 (***) are shown.

To determine the possible mechanism of root shortening in more details, we assayed the impact of *EXPA1*-overexpression on the longitudinal root zonation. Here, we used the cell morphology criteria as defined by Takatsuka *et al*. (2018). Our data show that while the number of cells in the TZ remained similar to those of WT, the number of cells in the RAM (the proximal meristem) was significantly reduced upon EXPA1 overexpression (Figure 9 E, F).

In contrast to previous reports, in our hands *expa1-1* (Pacifici *et al*., 2018), *expa1-2* (Ramakrishna *et al*., 2019) as well as our Dex-inducible amiRNA (amiEX1 lines), designed to downregulate *EXPA1* and closely related *EXPA14* and *EXPA15*, did not display any significant phenotype in terms of root or RAM size (Figure 9 - figure supplement 1). However, it should be noted here that the amiEX1 lines only reduced the *EXPA1* expression after Dex induction by app. 50%, *EXPA14* by 60% and *EXPA15* by 90% (tested by RT qPCR, data not shown).

Taking together, while we do not see any effect of *EXPA1* absence/downregulation on the root growth and/or RAM size, the overexpression of *EXPA1* results into reduction in a number of proliferating cells in the root meristem, thus slowing down the growth of the *Arabidopsis* root.

## DISCUSSION

### Is there a role for hormonal regulation over *EXPA1* in the root growth?

Recently, the role of *Arabidopsis* EXPA1 was described in the early stages of lateral root formation (Ramakrishna *et al*., 2019) and in the control of cell differentiation (expressed as a function of cell elongation) in the cells leaving meristematic zone of RAM (Pacifici *et al*., 2018). In the latter work, the authors proposed cytokinin-mediated upregulation of *EXPA1* and two genes encoding H^+^ATPases in the root TZ/EZ boundary and in the RAM, respectively, as a mechanism of cytokinin-induced cell differentiation. Pacifici *et al*. (2018) reported expansion of RAM in the *expa1-1* background compared to the WT. Furthermore, the authors claimed that the phenotype could be rescued in the presence of construct for translational fusion of *EXPA1* with GFP (pEXPA1::EXPA1:GFP), suggesting functionality of the construct even though no GFP signal could be detected. In line with more recent study (Ramakrishna *et al*., 2019), we did not observe any statistically significant change in the root length and/or RAM size in the *expa1-1*, CRISPR/Cas9 line *expa1-2* and our amiRNA lines. Nonetheless, it should be stressed here that *exp1-2* is hypomorphic allele (Ramakrishna *et al*., 2019) and our amiRNA lines are knock-down (not knock-out) lines. In contrast to Pacifici *et al*. (2018), we also did not observe the EXPA1:mCherry outside the columella/LRC in the root tip and the promoter activity (pEXPA1::nls:3xGFP) was only occasionally seen in the TZ/EZ boundary and in the elongated cells proximally to that. In accord with that, we did not detect any changes in the CW stiffness in the root TZ of *expa1-1*. Finally, cytokinins only moderately and transiently activated *EXPA1* transcription when assayed in the entire roots using RT qPCR. However, no statistically significant upregulation was detectable (using absolute fluorescence measurement) in the RAM using the pEXPA1::nls:3xGFP and pEXPA1::EXPA1:mCherry lines after 6h treatment with both BAP or NAA. These findings are suggesting that the cytokinin-mediated transcriptional regulation of *EXPA1* may take place in other parts of the root (e.g. cells surrounding LR primordia). Similarly to us, Pacifici *et al*. (2018) also did not see the cytokinin-dependent regulation in the columella/LRC and claim that *ARR1* mediates the cytokinin control over *EXPA1* expression specifically in the TZ/EZ boundary. Considering the *EXPA1* expression potentially taking place in the TZ/EZ boundary would represent negligible proportion of the entire expression of *EXPA1* in the root, its developmental importance is rather questionable and seems unlikely to be responsible for the observed (3-4 times) upregulation of *EXPA1* transcript in the *Arabidopsis* root. Based on our data, we do not exclude the role of EXPA1 in the control of RAM size, but probably in a concert with other EXPAs and dominantly in the columella/LRC (*vide infra*). The role of (cytokinin-regulated) auxin accumulation in the LRC in the control of RAM size has been proposed recently (Di Mambro *et al*., 2019). However, even in case of more distinct transcriptional regulation of *EXPA1* by NAA, we do not see any significant and consistent *EXPA1* upregulation in the LRC, both at the level of promoter activity or EXPA1 protein (data not shown), thus leaving the functional importance of cytokinin- and auxin-mediated regulation over *EXPA1* in the root tip rather unclear.

### EXPA localization and control of CW properties

In the previous studies, expansins were located to the CWs using immunolocalization techniques of fixed plant materials (Zhang and Hasenstein, 2000; Cosgrove *et al*., 2002; Balestrini *et al*., 2005). The transgenic lines carrying *EXPA* genes in a translational fusion with mCherry allowed observing localization of EXPA proteins, to our knowledge for the first time, in living plants. Interestingly, our data suggest that assayed *EXPAs* differ not only in the spatiotemporal specificity of expression, but their protein products reveal also distinct localization pattern in specific domains of the root apoplast. Firstly, we see localization of EXPA10 and EXPA14 dominantly in the longitudinal CWs of elongated root cells. This is resembling the situation observed in maize xylem, where the signal obtained after immunolocalization using anti-cucumber expansin antibody was homogenously distributed in the isodiametric (non-elongated) xylem cells, while located dominantly to the longitudinal CWs of elongated xylem (Zhang and Hasenstein, 2000). Secondly, EXPA10, EXPA14 and (partially) EXPA15 located in a punctuate pattern, spatially colocalizing with the three-cell boundaries, possibly surrounding the intercellular space. Plant as well as bacterial expansins were found to bind cellulose rather weakly (McQueen-Mason and Cosgrove, 1995), in case of bacterial EXLX1 via hydrophobic interactions (Georgelis *et al*., 2011). Much stronger affinity was observed between expansins and components of the CW matrix, including pectin and hemicelluloses (McQueen-Mason and Cosgrove, 1995; Georgelis *et al*., 2011). These and other evidence, e.g. the ability of expansins to mechanically weaken pure paper (McQueen-Mason and Cosgrove, 1994) led to a conclusion that expansins bind at the interface between cellulose microfibrils and polysaccharides in the CW matrix inducing the CW extension by reversibly disrupting the noncovalent bonds within this polymeric network (McQueen-Mason and Cosgrove, 1995). In the same study, the authors propose that an unknown minor structural component of the CW matrix might be responsible for expansin binding and action. In a more recent work using solid-state nuclear magnetic resonance (NMR) in a combination with differential isotopic labelling of expansin and polysaccharides, Wang et al. (2013) discovered that expansin binds highly specific cellulose domains enriched in xyloglucan, while the previously reported and more abundant binding to pectins doesn’t seem to relate to its activity. Our results imply existence of a factor determining specific localization of individual EXPAs in different CW compartments, particularly those revealing the specific “spotty” localization. The homogenous distribution of *EXPA1* throughout the CWs even outside its natural expression domain as seen in the pRPS5A>GR>EXPA1:mCherry line is suggesting that the specific localization pattern is not celltype specific, but rather encoded in the EXPAs amino acid sequence. The absence of colocalization of EXPA10 with calcofluor white staining in the fixed *Arabidopsis* root is implying that the factor could be a component of the CW matrix other than cellulose. The existence of the putative factor responsible for targeting subset of EXPAs into specific apoplast domains and its (molecular and/or biophysical) nature, however, remains elusive.

*Arabidopsis* genome contains 26 genes for α-expansins (Li *et al*., 2002), suggesting functional diversification within the subfamily. Specific expression and localization of *EXPA1*, EXPA10, EXPA 14 and EXPA15 together with differential hormonal sensitivity implies possible functional crosstalk among the individual expansins. Concert in their targeted action in the apoplastic continuum encapsulating the individual cells might result into the final vectorial change of CW expansion and highly coordinated cellular behaviour underlying root growth including its longitudinal zonation. Similar functional and spatiotemporal specificity including differential hormonal response and shoot cell growth-based zonation was described for *LeEXP2, LeEXP9* and *LeEXP18* in tomato (Caderas *et al*., 2000; Vogler *et al*., 2003). Using two independent approaches, the non-invasive Brillouin light scattering imaging and AFM we have shown that overexpression of *EXPA1*, homogenously distributed throughout the CW, results into increased CW stiffness in the root cells. That suggests that deregulating the tightly controlled equilibrium of specific expression and localization of individual EXPAs probably disturbs the naturally occurring strain/stress distribution within the growing *Arabidopsis* root that is reflected in the increased stiffness of the root cells and consequently root growth arrest. This might be analogous to the situation observed after misregulation of bipolar distribution of pectin methylesterase activity in the *Arabidopsis* hypocotyls. Peaucelle *et al*. (2015) demonstrated asymmetric loosening of longitudinal, as compared to transverse (anticlinal) walls just before the cell starts to elongate, even preceding the cortical microtubule reorientation, considered as a reporter of CW tensions (Hamant *et al*., 2019). It is achieved via asymmetric pectin de-methylesterification, as reliably shown via immunolabeling of low degree of homogalacturonan methylesterification in epidermal hypocotyl CWs using 2F4 antibody. As anticipated, manipulation of homogalacturonan demethylesterification through the inducible overexpression of the pectin methylesterase (PME5oe) or the PME inhibitor 3 (PMEI3oe) significantly increased or reduced 2F4 signal, respectively. That associated with reduction/increase of the overall cell stiffness in PME5oe/PMEI3oe plants. However, in both cases, the loss of asymmetry in the CW matrix composition and its biomechanical properties led to the similar effect, i.e. loss of cell expansion. Similar mechanism i.e. disturbing the tightly regulated spatial distribution of individual expansins might be the reason of contrasting effects of expansins overexpression, frequently associated with upregulated CW expansion, but sometimes leading to the opposite effect. i.e. cell growth inhibition (Caderas *et al*., 2000; Cho and Cosgrove, 2000, 2002; Choi *et al*., 2003; Vogler *et al*., 2003; Zenoni *et al*., 2011; Goh *et al*., 2014).

According to the loosening theory in a well-hydrated non-growing cell, the cell reaches osmotic equilibrium, with wall stresses counter-balancing the outward force of turgor pressure against the wall. However, growing CWs are loosened which refers to a shift or cut of a load-bearing part of the wall, relaxing tensile stress in the entire wall and simultaneously reducing cell turgor. As a result, water flows into the cell, elastically expanding the wall and restoring turgor and wall stress (Cosgrove, 2018b). CWs may become mechanically softer (meaning more easily deformed by mechanical force), but they do not necessarily result in an increase in wall relaxation and growth, e,g, lytic enzymes may soften CW but do not stimulate cell growth. On the other hand, α-expansins cause stress relaxation and prolonged enlargement of CWs, but they lack wall lytic activity and they do not soften the wall, as measured by tensile tests (Cosgrove 2018b, Wang and Cosgrove, 2020). An example of such observations was made by Wang and Cosgrove (2020) with pectin methylesterase (PME) that selectively softened the onion epidermal wall yet reduced expansin-mediated creep. After enzymatic de-esterification (without added calcium), the onion epidermal wall swelled and became softer, as assessed by nanoindentation (AFM) and tensile plasticity tests, yet exhibited reduced acid-induced creep. Accordingly, α-expansins were shown to act via different mechanism as compared to enzymes inducing CW creep via modifying CW matrix. Compared to CW loosening mediated by fungal endoglucanase Cel12A, expansins do induce CW loosening that is not associated with changes in the tensile stiffness (neither elastic nor plastic compliance), suggesting different way of action (Yuan *et al*., 2001). Another example is deesterification of homogalacturonan (HG) that is thought to stiffen pectin gels and primary CWs by increasing calcium crosslinking between HG chains. Contrary to this idea, recent studies (Braybrook and Peaucelle, 2013; Peaucelle *et al*., 2015) found that HG de-esterification correlated with reduced stiffness of living tissues, measured by surface indentation. The physical basis of such apparent wall softening is unclear, but possibly involves complex biological responses to HG modification. Indeed, feedback mechanisms and other factors regulating CW remodelling genes evoked by CW integrity pathway sensors often complicate the interpretation of CW mutant lines (Gigli-Bisceglia *et al*., 2018).

In terms of the methodological approach used, there are important differences between the longitudinal elastic modulus (M) measured by Brillouin light scattering and the Young’s modulus (E) measured via AFM (Prevedel *et al*., 2019). The BLS measured M is well known to be very sensitive to the level of hydration (Palombo *et al*., 2014; Wu *et al*., 2018; Androtis *et al*., 2019) and temperature (Berne and Pecora, 2000), and any comparisons have thus to be made under the same thermodynamic conditions and hydration levels. However, as these can be assumed to be similar between the different samples measured, variations between samples can be interpreted as being due to changes in the mechanical properties in the probed regime. The comparable trend of the AFM measured quasi-static Young’s Modulus and BLS measured MOC observed here, is consistent with observations in other diverse biological samples (e.g. Andriotis *et al*., 2019; Gouveia *et al*., 2029; Scarcelli *et al*., 2015) suggesting that here too the latter may serve as a proxy for stiffness. We note, however, that at very high hydration levels (much higher than in the system studied here), any relation between the two can be expected to break down (Wu *et al*., 2018).

One possible interpretation of the unexpected increase of CW stiffness upon *EXPA1* overexpression is the aforementioned disturbance of the coordinated equilibrium in the CW tensions across the *Arabidopsis* root and tissue-wide mechanical conflicts, shown to be an important morphogenic mechanism involved in the petal development in snapdragon (Rebocho *et al*., 2017). That might result to the general block in the CW extensibility, possibly potentiated via proposed mechanosensitive feedback loop (Uyttewaal *et al*., 2012), leading to absence of CW relaxation and thus increased stiffness. However, our large area mapping of CW stiffness using fluorescence emission–Brillouin imaging at small magnification implies colocalization of natural EXPA15 expression with regions of higher stiffness. This is implying a possible role for cell stiffening even in case of endogenous expansins. Nonetheless, the molecular/biophysical mechanism underlying this contra intuitive effect remains to be clarified.

### Conclusions

Based on our and others’ results (*vide supra*), we suggest that the tightly controlled spatiotemporal specificity of expansin expression in a combination with localisation of their protein products into distinct domains of plant extracellular matrix, together with hormone-regulated pH distribution within the root apoplast (Barbez *et al*., 2017), plays an important regulatory role controlling the root growth and development in *Arabidopsis*. Similar concept suggesting the role of regular (i.e. controlled) distribution of mechanically stiff regions in the extracellular matrix for the proper transcriptional regulation and actin-dependent cellular adhesion associated with stem cell fate determination was published in animal system (Yang *et al*., 2016).

The expansin-mediated regulation of the biomechanical CW properties seems to be associated with subsequent mechanoperception-mediated feedback loop, leading to genome-wide changes in the expression profiles of individual cells, probably in a cell type-specific context (Ilias *et al*., 2019). However, our results on the short-term induction of *EXPA1* expression associated with prompt increase in the CW stiffness imply that the non-transcriptional regulation will be an important mechanism underlying EXPA(1)-controlled CW biomechanics and root growth.

Upregulated *EXPA1* associated with CW stiffness seems to downregulate root growth via downregulating RAM size. This is suggesting a mechanism connecting biomechanical CW properties with the control over cell division in the RAM. Whether the mechanism includes the CW integrity signalling, previously shown to control cell division in the RAM in a response to the inhibition of cellulose biosynthesis (Gigli-Bisceglia *et al*., 2018), remains to be identified.

## ACKNOWLEDGEMENT

M.S. has received funding from the European Union’s Horizon 2020 research and innovation programme under the Marie Skłodowska-Curie co-financed by the South Moravian Region under grant agreement No. 665860. This study reflects only the author’s view and the EU is not responsible for any use that may be made of the information it contains. The work was further supported by the Ministry of Education, Youth and Sports of CR from European Regional Development Fund-Project “Centre for Experimental Plant Biology”: No. CZ.02.1.01/0.0/0.0/16_019/0000738, LTAUSA18161 and LTC19047 and Czech Science Foundation (19-24753S). K.E. acknowledges support from the City of Vienna and the Austrian Ministry of Science (Vision 2020). The work of E.V.Z. and E.V.U. was supported by the Russian Science Foundation (20-14-00140) and the Russian State Budget (0324-2019-0040). We acknowledge Plant Sciences and Cellular Imaging CFs at CEITEC MU supported by MEYS CR (LM2018129). RIAT-CZ project (ATCZ40) funded via Interreg V-A Austria – Czech Republic is gratefully acknowledged for the financial support of the measurements at the Vienna Biocenter CF Advanced Microscopy. We thank to Ive De Smet for his kind donation of the pEXP1::nls:3xGFP, Alexis Maizel for CRISPR/Cas9 *exp1-2 Arabidopsis* lines and Victoria Mironova for critical reading of the manuscript.

## MATERIAL AND METHODS

### Promoter analysis

2500 bp regions upstream of the transcription start site and entire 5’UTRs were used for prediction of TF binding regions in gene promoters. The TAIR10 version of the *A. thaliana* genome (https://www.arabidopsis.org/download_files/Genes/TAIR10_genome_release/TAIR10_chromosome_files/TAIR10_chr_all.fas) was used for the analyses. *A. thaliana* genome annotation data were retrieved from Araport11 (https://www.arabidopsis.org/download_files/Genes/Araport11_genome_release/Araport11_GFF3_genes_transposons.201606.gff.gz). To identify potential B-ARR binding regions, two sets of publicly available ChIP-seq data we used. First on ARR1,10,12 binding in 3-day old seedlings of Ypet-tagged B-ARRs lines treated with 10 uM BAP or mock treated for 4h (Xie *et al*., 2018). Second on ARR10 binding in two to 3-week old seedlings of 35S::ARR10:GFP lines treated with 5 uM BAP or mock treated for 30 minutes (Zubo *et al*., 2017). To identify potential ARF binding regions we used DAP-seq data for ARF2 and ARF5 (O’Malley *et al*., 2016). The corresponding processed data were retrieved from Gene Expression Omnibus database (https://www.ncbi.nlm.nih.gov/geo/), the Quickload server for the Integrated Genome Browser (IGB) (bioviz.org, Freese *et al*., 2016) and Plant Cistrome Database (http://neomorph.salk.edu/PlantCistromeDB), respectively.

### Quantitative real-time transcript profiling (RT qPCR)

Total RNA was extracted from 7-day old wild-type *Arabidopsis thaliana* (ecotype Columbia-0) seedlings treated with either 5 μM BAP or 5 μM NAA for 0.5h, 1 h, 2h, 4h and non-treated seedlings as controls. First-strand cDNA was synthesized from total RNA using SuperScript III Reverse Transcriptase (Thermo Fisher Scientific). RT qPCR was performed on cDNAs, with primers spanning an intron summarized in Table 1: for *EXPA1* (*At1g69530*, P1 and P2), *EXPA10* (*At1g26770*, P3 and P4), *EXPA14* (*At5g56320*, P5 and P6) and *EXPA15* (*At2g03090*, P7 and P8). The transcript abundance of *EXPAs*, relative to constitutively expressed normalizer gene, UBQ10 (*At4g05320*, P9 and P10), was quantified, using the 2(-Delta Delta C(T)) method (Livak and Schmittgen, 2001) and calibrated to expression at 0h (non-treated). Real-time quantification was performed in Rotor-Gene Q 72-slots using the Rotor-Gene Q Series Software (QIAGEN). PCR conditions were: 95°C for 7 min, one cycle; 15 s at 95°C, 30 s at 56°C, 30 s at 72°C, 40 cycles.

**Table 1:**
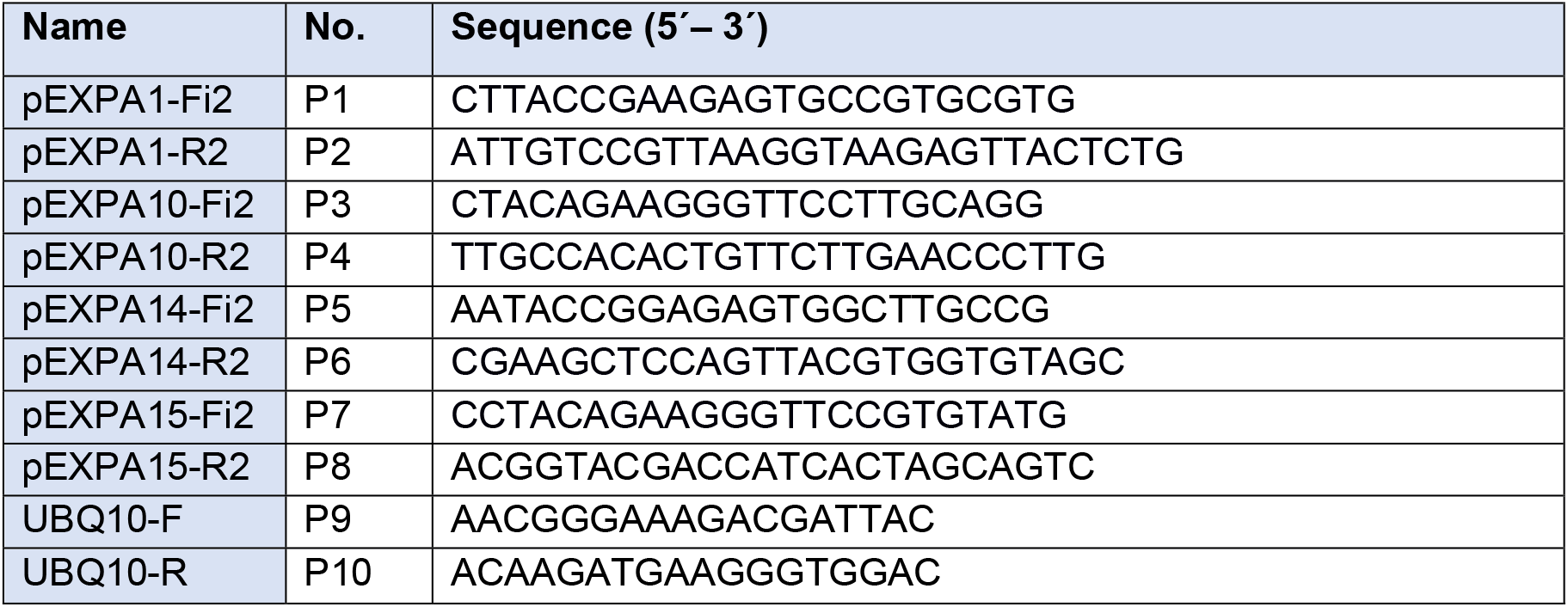
List of primers used for RT qPCR.

Reactions with no cDNA monitored for the presence of primer dimers and no reverse transcriptase controls were included for each cDNA sample. PCRs were carried out in triplicate and mean values determined.

### Cloning and plant transformation

Standard molecular techniques as described by Ausubel *et al*., (1999) were used. To clone the translational fusions of expansins with mCherry, firstly, an intermediate clone (pZEO-mCherryT35S) that contains unique restriction sites PacI and SnaBI as well as a flexible linker in front of mCherry sequence, was created as follows. Two DNA fragments were generated by polymerase chain reaction (PCR) using Herculase II Fusion DNA polymerase (Agilent Technologies), primers P11 + P12 and P13 + P14 and plasmids pUCAP-pGEL3::spmCherry-pATrpC-BAR (Samalova *et al*., 2017) and pOpIn2 (Samalova *et al*., 2019) as templates for mCherry and a polyadenylation signal (T35S) respectively. The fragments were joined together by overlapping PCR using P11 and P14. The final product was cloned by BP reaction into attB1 and attB2 sites of GATEWAY™ compatible plasmid pDONOR/Zeo and confirmed by sequencing. Secondly, individual promoter sequences together with *EXPA* coding sequences (but without a stop codon) were amplified from genomic DNA using primers P15 + P16 (pEXPA1::EXPA1), P17 + P18 (pEXPA10::EXPA10), P19 + P20 (pEXPA14::EXPA14) and P21 + P22 (pEXPA15::EXPA15). The products were digested with either *Sna*BI (pEXPA1::EXPA1 and pEXPA15::EXPA15), *Pac*I (pEXPA10::EXPA10) or both *Pac*I/*Sna*BI (pEXPA14::EXPA14) and cloned into the same sites of pZEO-mCherryT35S and confirmed by sequencing. Finally, the pEXPA::EXPA:mCherryT35S fusions were re-cloned by LR reaction into attR1 and attR2 sites of pFAST-G01 vector (Shimada *et al*., 2010).

To overexpress *EXPA1* and *EXPA1:mCherry* fusion, the dexamethasone (Dex) inducible pOp6/LhGR system (Craft *et al*., 2005; Samalova *et al*., 2005) was used. Firstly, we PCR-amplified EXPA1 using P23 + P24 and EXPA1:mCherry using P23 + P25 sequences from the pZEO-pEXPA1::EXPA1:mCherryT35S vector generated above, cloned into the pDONOR/Zeo vector and confirmed by sequencing. Secondly, using GATAWAY™ cloning strategy described above we re-cloned the EXA1 and EXPA1;mCherry sequences into a pOpIn2-RPS5A plasmid (Samalova *et al*., 2019) that drives the LhGR activator under the constitute AtRPS5A promoter (Weijers *et al*., 2001).

*Arabidopsis* thaliana (ecotype Columbia-0) was transformed using the floral dip method (Clough and Bent, 1998) and the transgenic plants were selected on Murashige and Skoog (MS) medium (Murashige and Skoog, 1962) containing 15 μg/ml hygromycin for the pFAST-G01 vectors and 10 μg/ml phosphinothricin for the pOpIn2 vectors.

**Table 2:**
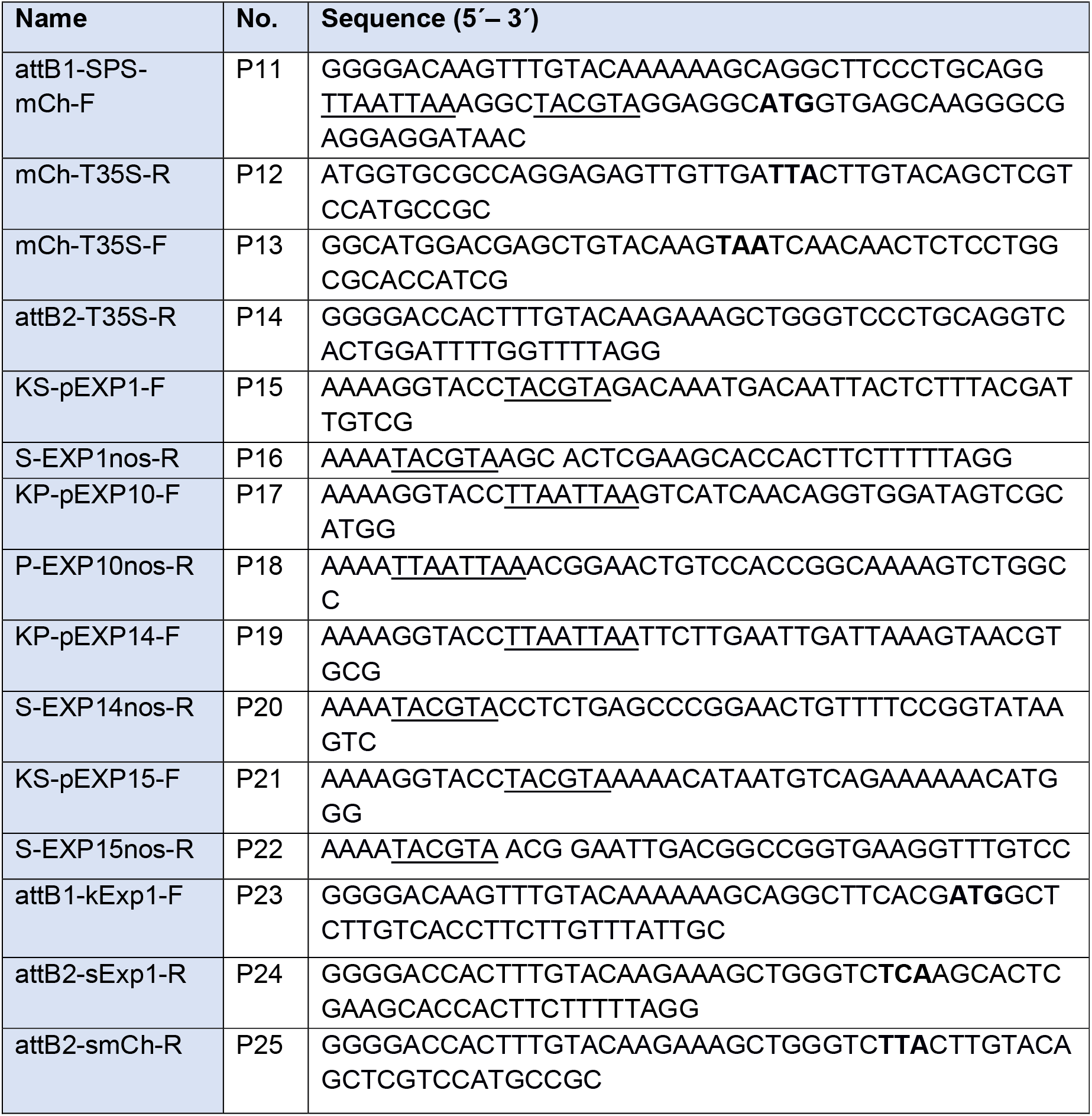
**List of primers used for expansin cloning** – underlined unique restriction sites used for cloning, **bold** start and stop codons.

### Plant growth condition and dexamethasone (Dex) induction

Standard MS medium supplemented with 1.5% sucrose and 0.8% plant agar (Duchefa), pH 5.8 adjusted by KOH or pH 4 adjusted by H_2_SO_4_ was used. Plants were cultivated in growth chambers under long day conditions (16 h light/ 8 h dark) at 21°C in Petri dishes or in soil, with a light intensity of 150 μM.m^-2^.s^-1^ and 40% relative humidity. Unless otherwise stated induction was performed by adding 20 μM Dex into the media as described in Samalova *et al*. (2019). DMSO at the same concentration was used as a control.

### Confocal laser scanning microscopy (CLSM) and image analysis

To localise EXPA:mCherry fusions Zeiss LSM 880 laser-scanning microscope was used. mCherry fluorescence was detected at 580-650 nm with 561-nm HeNe laser excitation and eGFP at 490-550 nm with a 488-nm Argon laser line. Z-stack projections are shown as maximum intensity. Quantification of the fluorescence was done using CellProfiler (mCherry) and Imaris (nls:3xGFP) softwares. To measure the size of RAM, 7-day old *Arabidopsis* seedlings were stained with propidium iodide at concentration 30 μg/ml for 5 min, scanned at 590-650 nm with 488-nm excitation and measured using the ZEN 3.0 software. The roots were imaged using the C-Apochromat 40x/1.2 water corrected objective lens or Plan-Apochromat 25x/0.8 immersion corrected.

### Brillouin light scattering (BLS) microscopy

Brillouin microscopy was performed using a homebuilt Brillouin confocal microscope described in Elsayad *et al*., 2016. Excitation was via a single-mode 532nm laser (Torus, Laser Quantum, DE). A dual cross-dispersion Virtual Imaged Phase Array (VIPA) spectrometer (Scarcelli *et al*., 2015) with a Lyott Stop (Edrei *et al*., 2017) was employed for measuring the Brillouin Light Scattering spectra. The spectral projection was measured on a cooled EM CCD camera (ImageEMX II, Hamamatsu, JP). The spectrometer was coupled to an inverted microscope frame (IX73, Olympus, JP) via a physical pinhole with an effective size of 1 Airy Unit to assure optimum confocal detection. After the pinhole a dichroic mirror was used to outcouple light with wavelengths longer than 536nm to a fluorescence spectrometer (Ocean Optics QE Pro, USA) to detect the fluorescence signal assuring pixel-to-pixel correlation with the measured Brillouin spectra. To acquire Brillouin maps, samples were scanned in x,y &/or z using either a 3-axis long-range Piezostage (Physik Instrumente, DE) or a motor stage (ASI, USA), both mounted on top of the inverted microscope frame. Light could also be coupled out through a second port on the microscope frame using a long-pass filter (AHF, DE) and tube lens to a compact sCMOS camera (Thorlabs, DE) allowing us to locate samples and regions of interest (in wide-field transmitted light conditions when illuminating sample from the top with a Halogen lamp) as well as monitor the position being probed during scanning.

All hardware was controlled using LabView (National Instruments, USA) based software developed by the company THATec (DE) especially for our microscope. The 16bit depth spectral projection image for each position in a spatial scan was exported from the native THATec format into Matlab (Mathworks, DE), where a custom written code was used for analysis. This code (see also Elsayad *et al*., 2016) used two calibration spectra (of triple distilled water and spectroscopic grade ethyl alcohol) measured before and after each set of scans. These were used for registration of the spectral projection onto a frequency scale, based on the calculated disperion for a dual-VIPA setup in the paraxial approximation regime (Xiao *et al*., 2014). The alignment of the spectrometer was such that maximal energy was transferred into a single diffraction order. Due to the spatial masking of the elastic scattering peaks at the two intermediate imaging planes in the spectrometer, the spectral projection consisted of only two inelastic scattering peaks corresponding to the so-called Brillouin Stokes and anti-Stokes scattering peaks.

All data analysis was performed in Matlab (Mathworks, DE) using custom written scripts (Elsayad *et al*., 2016). Spectral phasor analysis (Elsayad, 2019) was used to obtain initial parameter estimates for peak positions and widths which were subsequently inserted into a non-linear least squares fitting algorithm that fitted two broadened Lorentzian functions (Voigt functions) to obtain the two peak positions, from which the Brillouin frequency shift could be obtained. The BLS spectra was also deconvolved in phasor space using a response function obtained from measuring the attenuated Rayleigh scattering inside the respective samples (by opening the spatial masks).

For all scans the laser power at the sample was between 1-5 mW, and the dwell time per point, which was also the acquisition time of each spectra, was 100ms. Cells were observed (by transmitted-light widefield illumination) to appear healthy and unperturbed after experiments, suggesting the BLS measurements had no negative or phototoxic effects. A 1.4 NA objective was used for excitation and detection (back-scattering geometry). As such a broad range of scattering wavevectors is probed and one effectively probes directionally averaged elastic moduli. As a direct consequence of probing a broad spectrum of wavevectors, the Brillouin spectra is broadened as predicted from the momentum-energy conservation equations describing the scattering processes. The so-called Brillouin scattering peak position is however not noticeably modified to within experimental uncertainties, as was verified by reducing the numerical aperture of excitation and detection on the studied samples using an iris in the beam path.

Roots of 7-day old *Arabidopsis* seedlings were scanned at the early EZ, the size of the scan was 25 um x 25 um (50 x 50 pixels), step size typically 500 nm (or 250 nm for larger scans) using the piezo-stage.

### Refractive index tomography

Refractive index tomograms were acquired on a holotomographic microscope with rotational scanning 3D Cell Explorer (Nanolive SA, Lausanne, Switzerland) with Nikon BE Plan 60x NA 0.8. The size of acquired tomogram was 93.1×93.1×35.7 μm (xyz). Samples were measured in water (reference refractive index 1.330). Software Steve 1.6.3496 (Nanolive SA) was used for image acquisition. Subsequent image analysis was performed in ImageJ 1.52q (NIH, USA) on a max projection of tomography data. Following parameters were extracted: mean refractive index at cell wall of longitudinal and transverse axes of cell.

### Atomic force microscopy (AFM)

Roots of 7-day old *Arabidopsis* seedlings were immobilized on glass slides and surrounded by stiff agarose. Approximate early EZ was defined based on the visual landmark observed through a bright field microscope. In order to extract the mechanical properties of only the outer cell wall, the maximum indentation force was set to 60 nN to archive a maximum indentation of no more than 80 nm. The cantilever used was “Nano World” (Nanosensors Headquarters, Neuchâtel, Switzerland) SD-R150-T3L450B tips with a spring constant of 0.15–1.83N/m (the one used was estimated to be 0.781 N/m) with silicon point probe tips of a 150-nm radius.

All force spectroscopy experiments were performed as previously described (Feng *et al*., 2018; Peaucelle, 2014; Peaucelle *et al*., 2015). Briefly, stiffness of samples was determined as follows: an AFM cantilever loaded with a spherical tip was used to indent the sample over a 60 × 100 μm square area, within the area 60 × 100 measurements were made resulting in 6000 force-indentation experiments; each force-indentation experiment was treated with a Hertzian indentation model to extract the apparent Young’s modulus (E_A_); each pixel in a stiffness map represents the apparent Young’s modulus from one force-indentation point. The E_A_ was calculated using the JPK Data Processing software (ver. Spm - 4.0.23, JPK Instruments AG, Germany), which allows for a more standardized analysis than the estimation of the E_A_ using a standard Hertzian contact model (Peaucelle, 2014; Peaucelle *et al*., 2015). Only the retraction curve was used in our analyses as is typically the case in nano-indentation experiments. A Poisson ratio of 0.5 was assumed for the material. Range distribution of E_A_ from 0.2 MPa to 3 MPa in 1-MPa binned groups was calculated using MATLAB.

### Statistical analysis

For statistical analyses simple ANOVA and post-hoc Tukey test was used. For pairwise comparisons in repeated experiments, mixed model ANOVA using random effects for the different experiments was used with Tukey test as a post-hoc test. In case of non-normal count data (e.g No. of cells) a Poisson mixed model was used to identify differences between genotypes. For the implementation of the mixed models the lme4 package in R was used (Bates *et al*., 2015).

## SUPPLEMENTARY FIGURES AND FIGURE LEGENDS

**Figure 1 – Figure Supplement 1:**
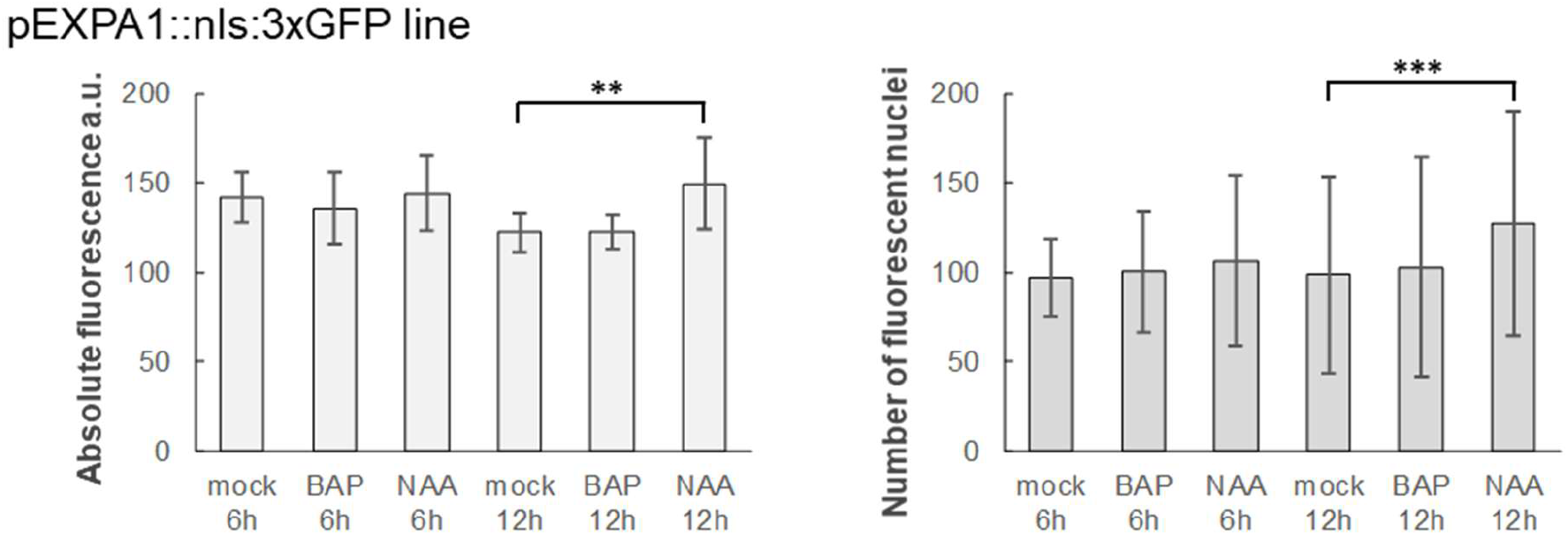
*EXPA1* is not inducible by cytokinins in the root tip. 7-day old *Arabidopsis* seedlings of pEXPA1::nls:3xGFP line were treated with 5 μM BAP and 5 μM NAA for 6h and 12h. Absolute fluorescence (left) and a number of fluorescent nuclei (right) was quantified from 3D confocal z-stacks using IMARIS software. 8-10 seedlings were used in each category, average and SD were calculated and displayed in the graphs. Statistically significant differences at alpha 0.01 (**) and 0.001 (***) are shown.

**Figure 1 – Figure Supplement 2:**
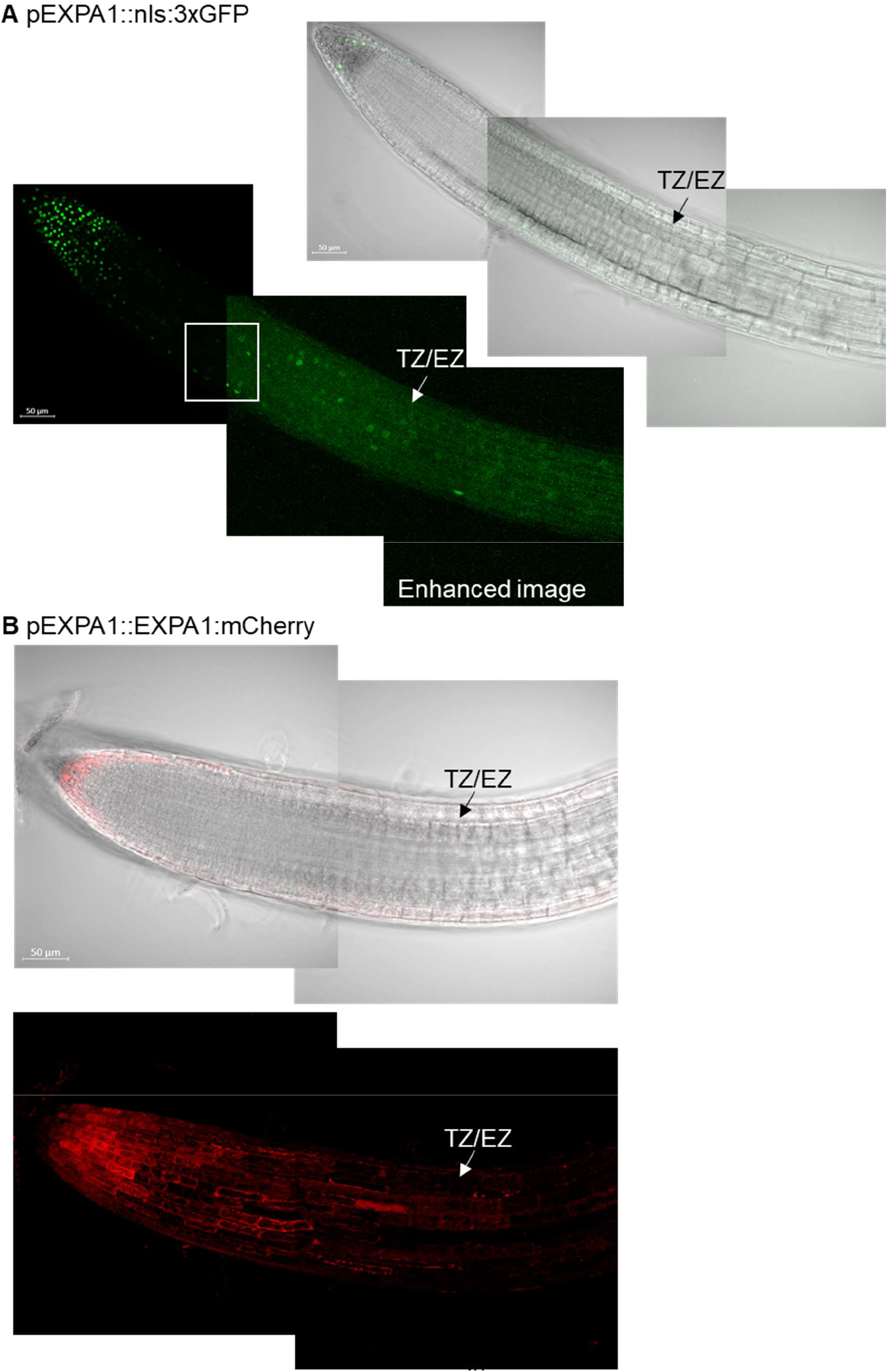
*EXPA1* activity is only occasionally detectable in the root TZ while EXPA1 signal is delimited to the columella/lateral root cap. Z-stack projections and transmitted-light micrographs shown as a single optical section of 7-day old Arabidopsrs seedlings of **(A)** pEXPA1::nls:3xGFP and **(B)** pEXPA1::EXPA1:mCherry lines. The TZ/EZ boundary is depicted (arrow). The white square marks the same root area visualised without and with image enhancement done using the CLSM Zen 3.0 software.

**Figure 2 – Figure Supplement 1:**
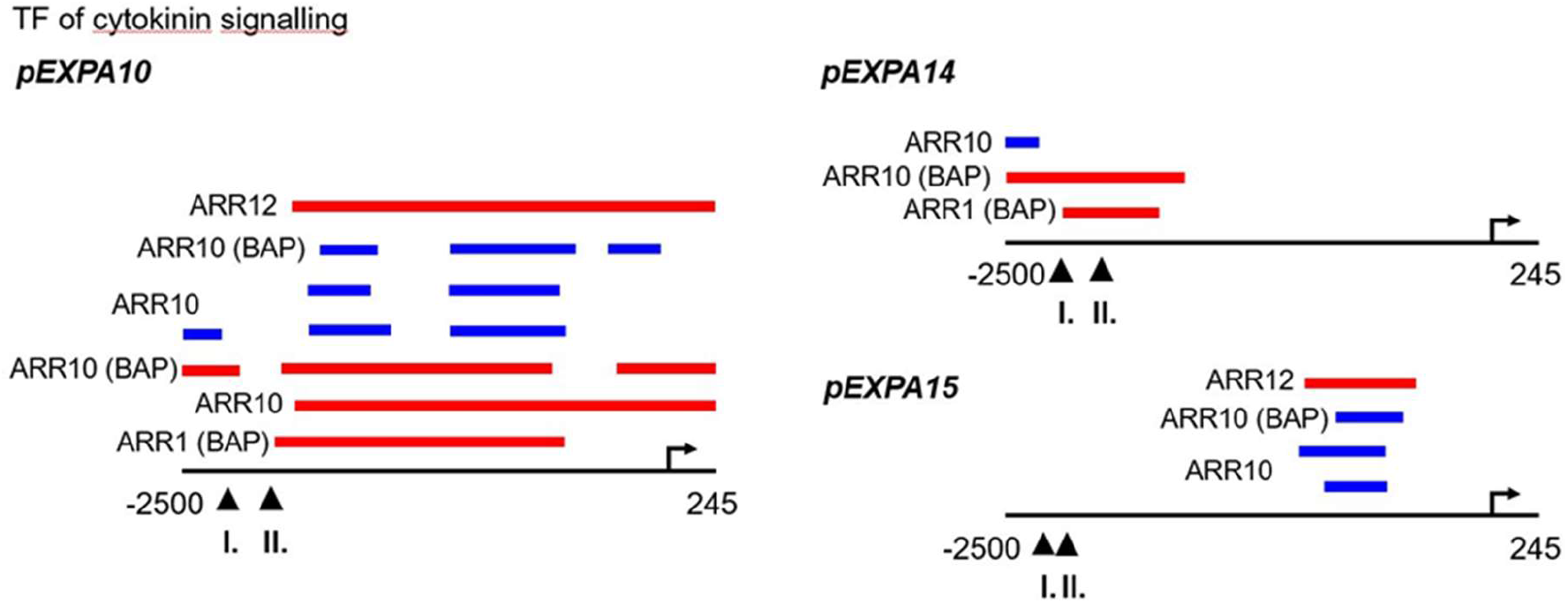
Cytokinin-responsive transcription factors bind to *EXPA* promoters. Promoter analysis of *EXPA10, EXPA14* and *EXPA15* identifies ChIP-seq derived binding events for transcription factors (type-B ARABIDOPSIS RESPONSE REGULATORS, ARRs) involved in the cytokinin signalling pathway. Red, blue and green colours depict the peaks from Xie *et al*., 2018, Zubo *et al*., 2017 and O’Malley *et al*., 2016, respectively. The coordinates are represented relative to the transcription start site marked by the arrow. The arrowheads indicate the 5’-end of each promoter as used in (I.) this publication and (II). Pacifici *et al*., 2018.

**Figure 3 – Figure Supplement 1:**
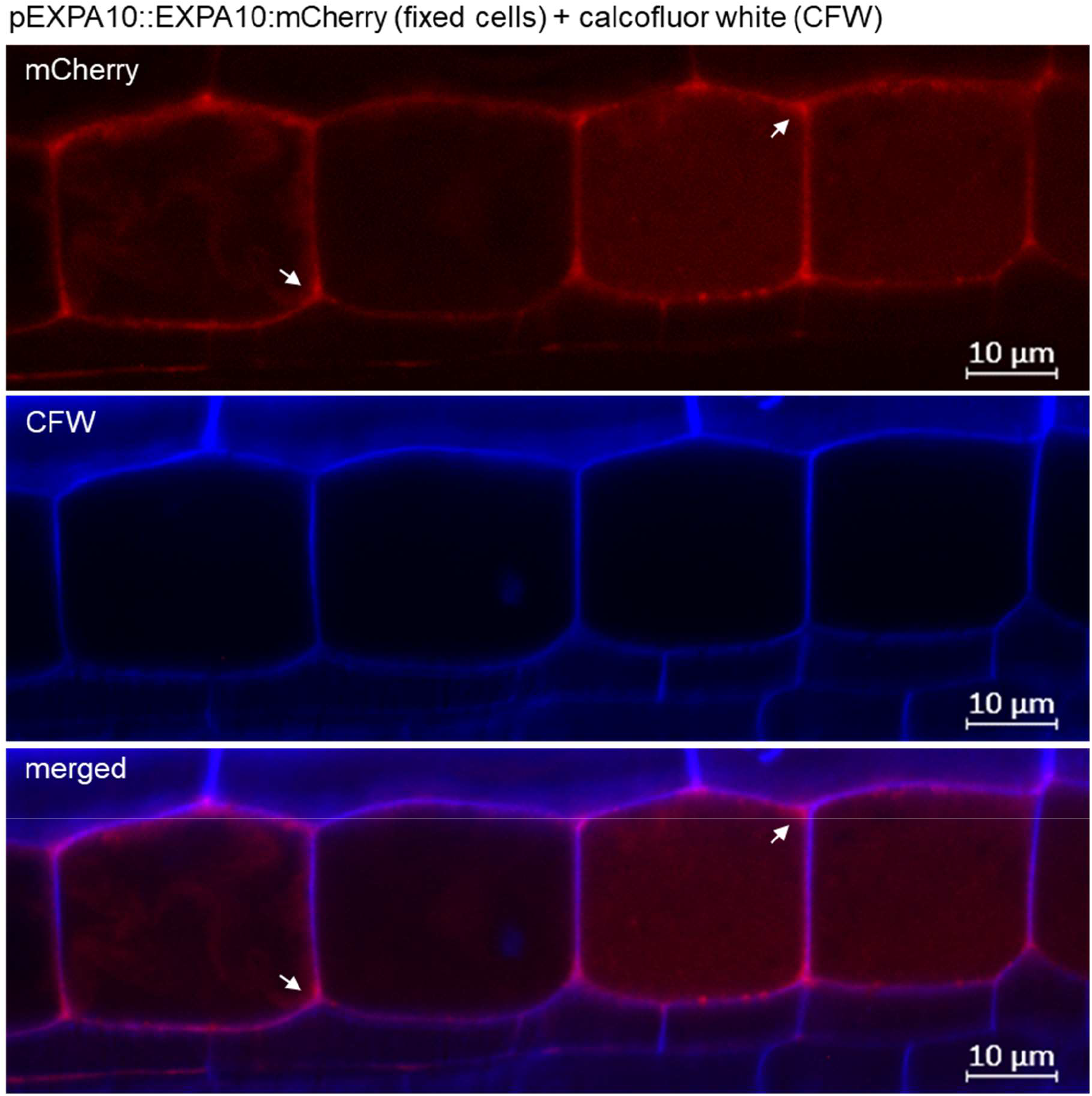
EXPA10 does not completely colocalize with cellulose in fixed cell walls. Z-stack projections of a pEXPA10::EXPA10:mCherry line (red) fixed, cleared and stained with calcofluor white (CFW, blue) according to a protocol in Ursache *et al*., 2018. The arrowheads point to positions where the EXPA10:mCherry does not co-localise with cellulose deposition as labelled by CFW. Scale bar 10 μm.

**Figure 3 – Figure Supplement 2:**
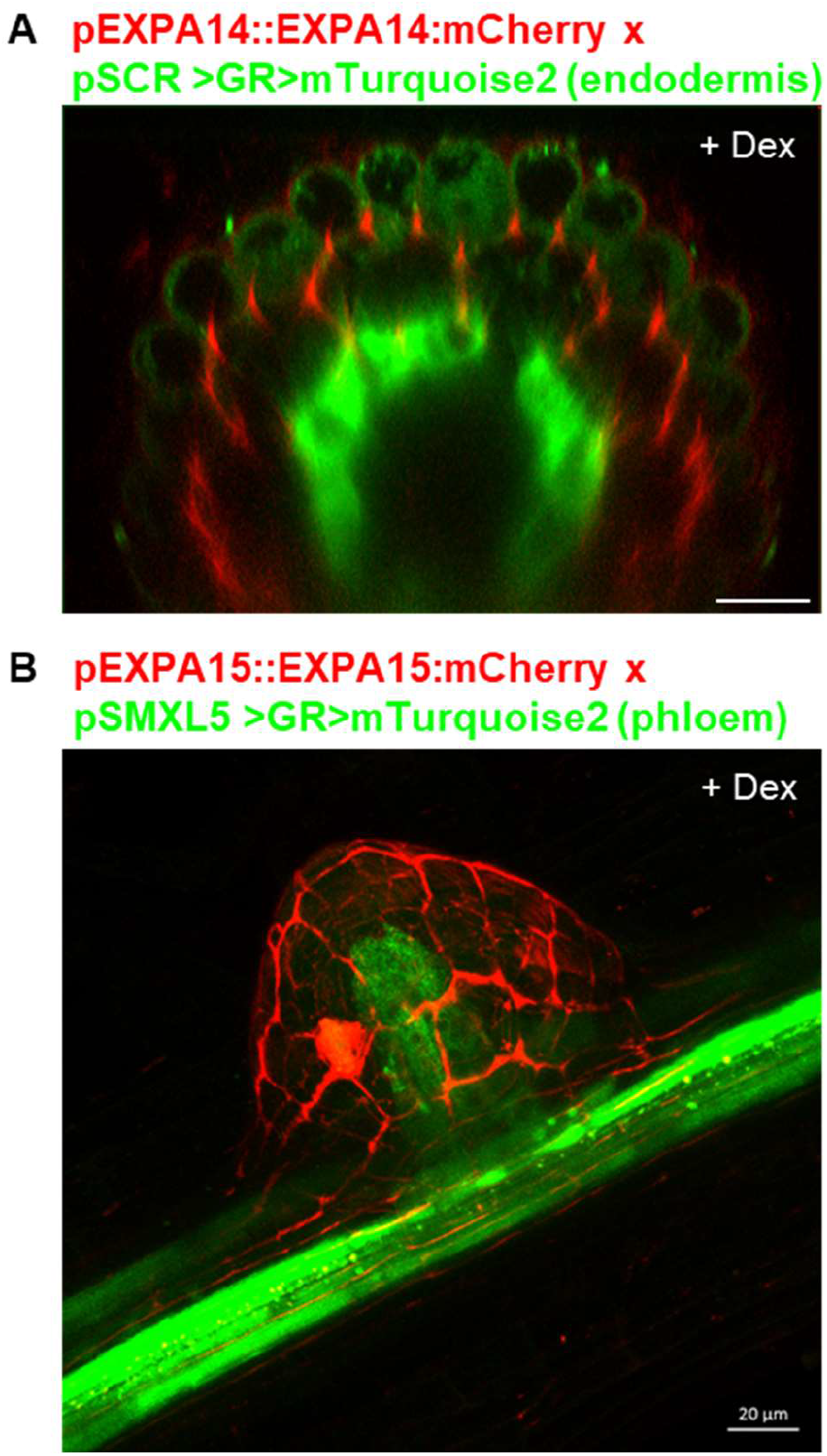
Cell-type specific localization of EXPA14 and EXPA15. **(A)** Double hemizygous F1 progeny of a cross between *pEXPA14::EXPA14:mCherry* and endodermis-specific *pSCR>GR>mTurquoise2* (Schurholz *et al*., 2018). In the transversal (xz) optical section of a primary root, the EXPA14 (red) localizes to cortex, endodermis and background autofluorescence in epidermis is in green. **(B)** Double hemizygous F1 progeny of a cross between *pEXPA15::EXPA15:mCherry* and phloem-specific *pSMXL5>GR>mTurquoise2* (Schurholz *et al*., 2018)In the z-stack projection of an emerging lateral root EXPA15 (red) localizes to the CW of epidermis, phloem is in green. The 7-day old Arabidopsis seedlings were grown on MS media +Dex to induce the mTurquoise2 ER-specific expression. Scale bars correspond to 20 μm.

**Figure 3 – Figure Supplement 3:**
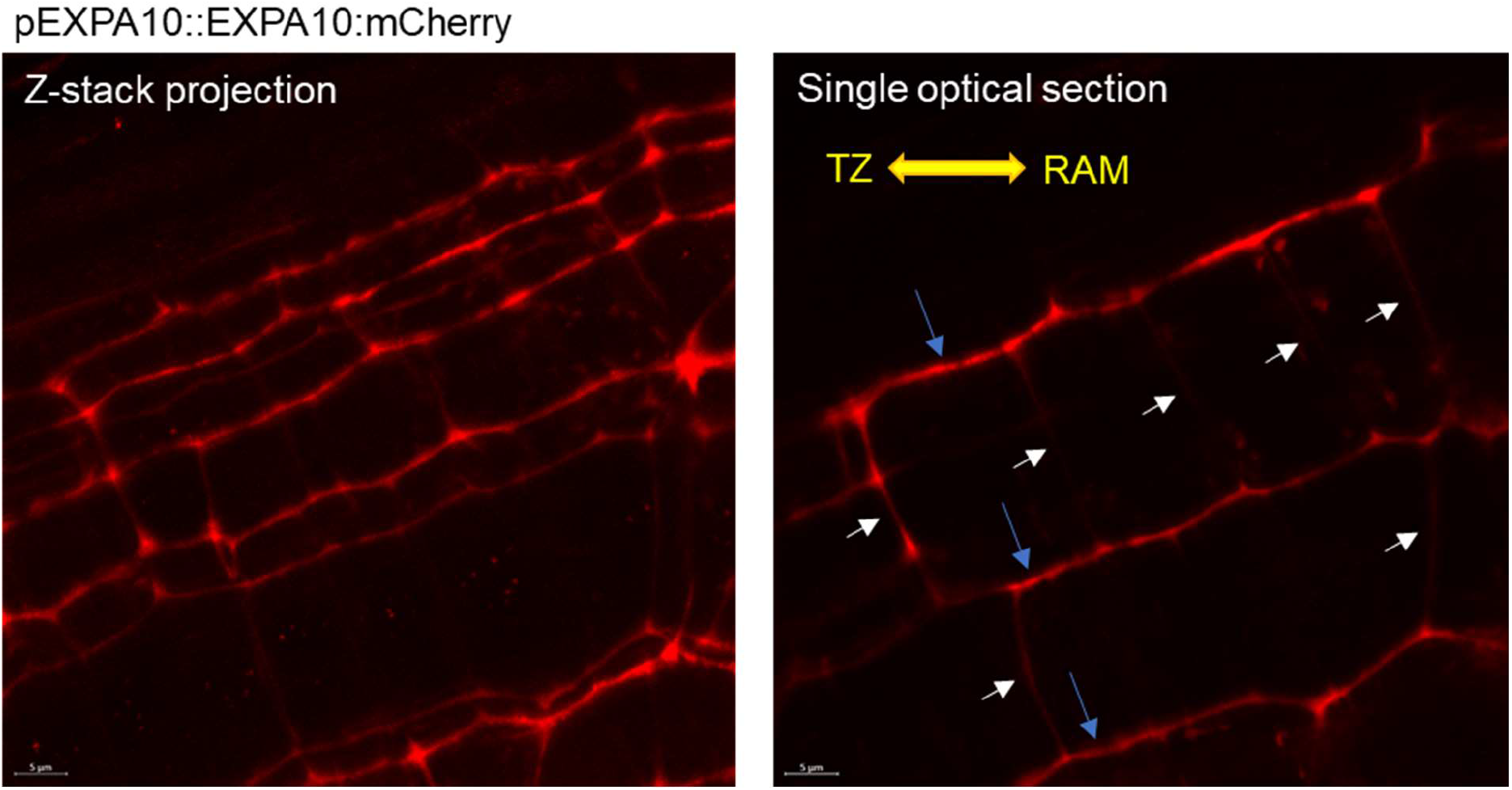
EXPA10 localizes predominantly to the longitudinal cell walls. A z-stack projection (, left) and a single optical section (right) of a 7-day old *Arabidopsis* root of pEXPA10::EXPA10:mCherry line imagined using the Airyscan detector of Zeiss 880 CLSM. Blue arrows point to longitudinal while white arrows to (less visible due to lower EXPA10 signal intensity) transversal CWs of individual cells; the root orientation is indicated by the yellow double arrows. Scale bar 5 μm.

**Figure 4 – Figure Supplement 1:**
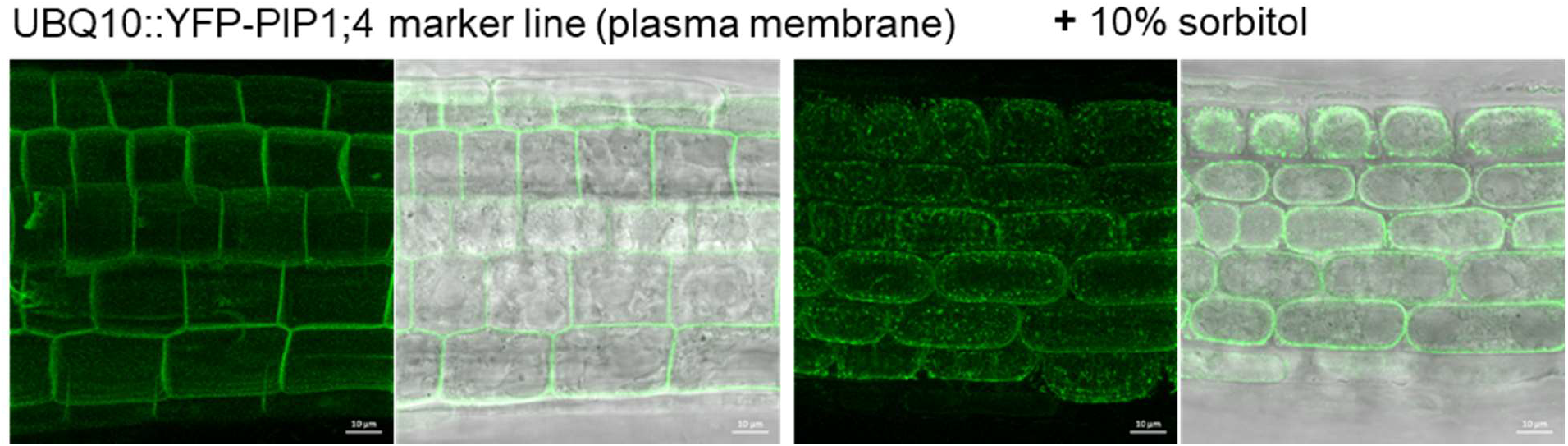
Confocal imaging of a plasma membrane marker line UBQ10::YFP-PIP1;4 before and after plasmolysis. Z-stack projections and transmitted-light micrographs shown as a single optical section of 7-day old *Arabidopsis* seedlings of a UBQ10::YFP-PIP1;4 line (von Wangenheim *et al*., 2016) labelling plasma membrane imaged before and after treatment with 10% sorbitol for 10 min. Scale bar 10 μm.

**Figure 5 – Figure Supplement 1:**
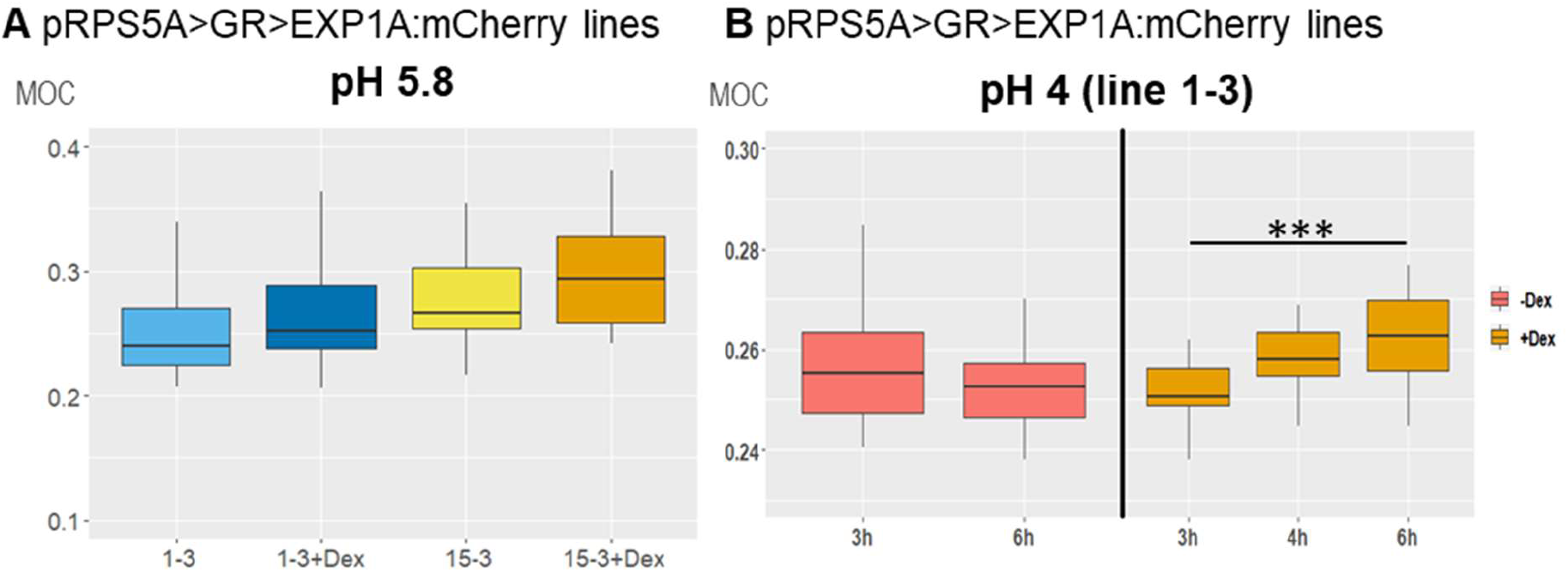
Low-level *EXP1A:mCherry* overexpression reveals only weak cell wall stiffening determined using Brillouin light scattering microscopy. Mechano-Optical Contrast (MOC) was determined in roots of 7-day old *Arabidopsis EXPA1:mCherry* overexpressing seedlings pRPS5A>GR>EXPA1:mCherry (lines 1-3 and 15-3) grown on MS media **(A)** +/- Dex pH 5.8 or **(B)** induced in liquid MS media pH 4 for 3h - 6h; DMSO was used in -Dex treatments. Medians shown are from at least 4 seedlings and 10 measurements in each category. Statistically significant differences at alpha 0.001 (***) are shown.

**Figure 5 – Figure Supplement 2:**
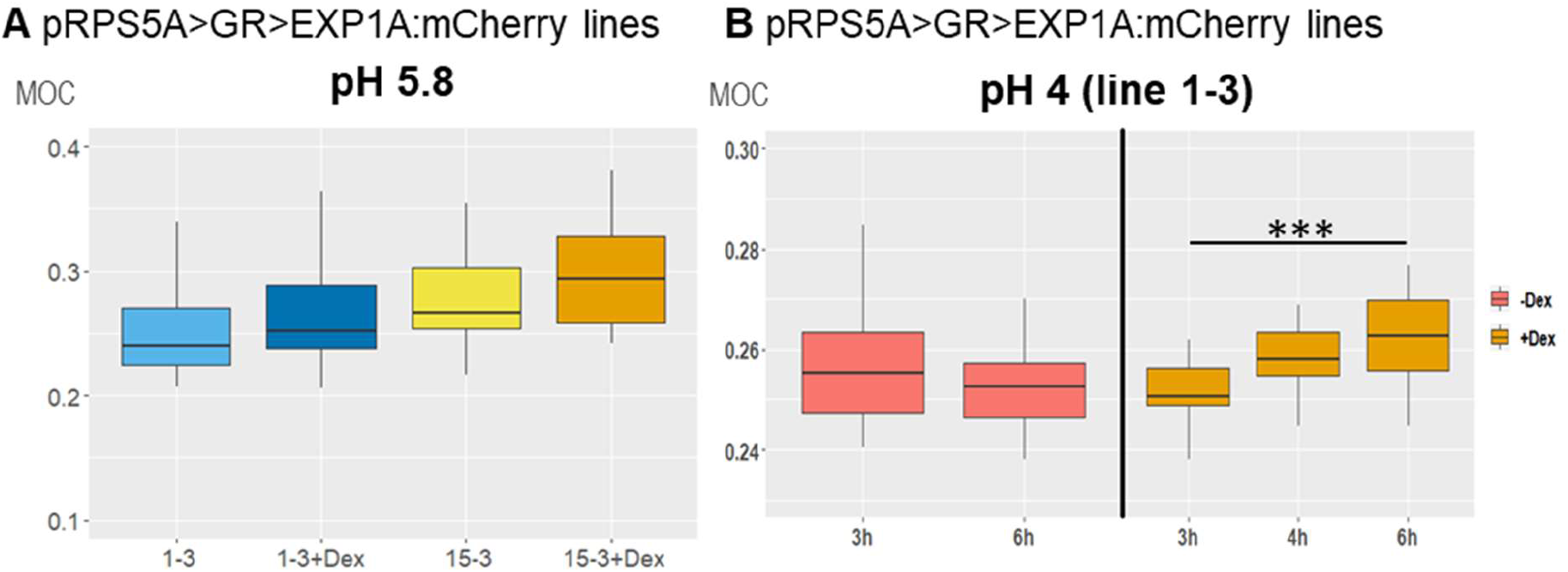
Quantitative real-time PCR of Dex-induced levels of expression of *EXP1A:mCherry* and *EXPA1* in selected lines. Relative *EXPA1* expression levels in independent single-copy T3 homozygous lines of **(A)** pRPS5A>GR>EXP1A:mCherry (1-3, 3-2, 9-1, 15-3, 16-1 and 18-1) and **(B)** pRPS5A>GR>EXP1A (4-1, 5-4, 6-3, 8-4, 12-4 and 17-1) seedlings grown on MS media +/- Dex for 7 days. The transcript abundance of *EXPA1* is double normalized to *UBQ10* and WT controls. The experiment was done once with 3 technical replicas. Note the logarithmic scale of the *EXPA1* relative expression.

**Figure 5 – Figure Supplement 3:**
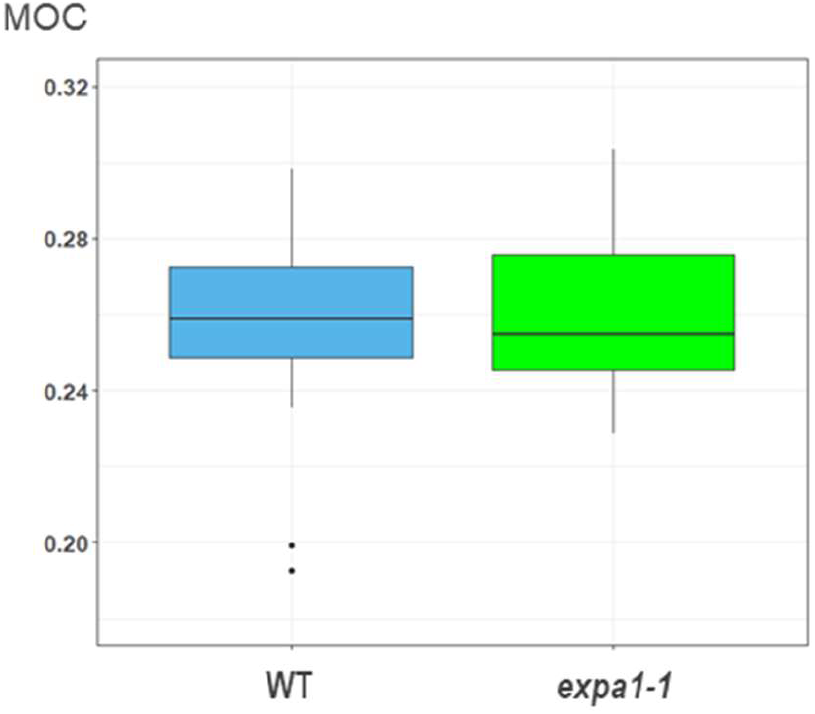
Knock-out *expa1-1* does not show changes in cell wall stiffening in the TZ determined using Brillouin light scattering microscopy. Mechano-Optical Contrast (MOC) was determined in roots of 7-day old *Arabidopsis* WT and *expa1-1* knock-out line. Medians shown are from at least 4 seedlings and 12 measurements in each category. There are no statistically significant differences.

**Figure 9 – Figure Supplement 1:**
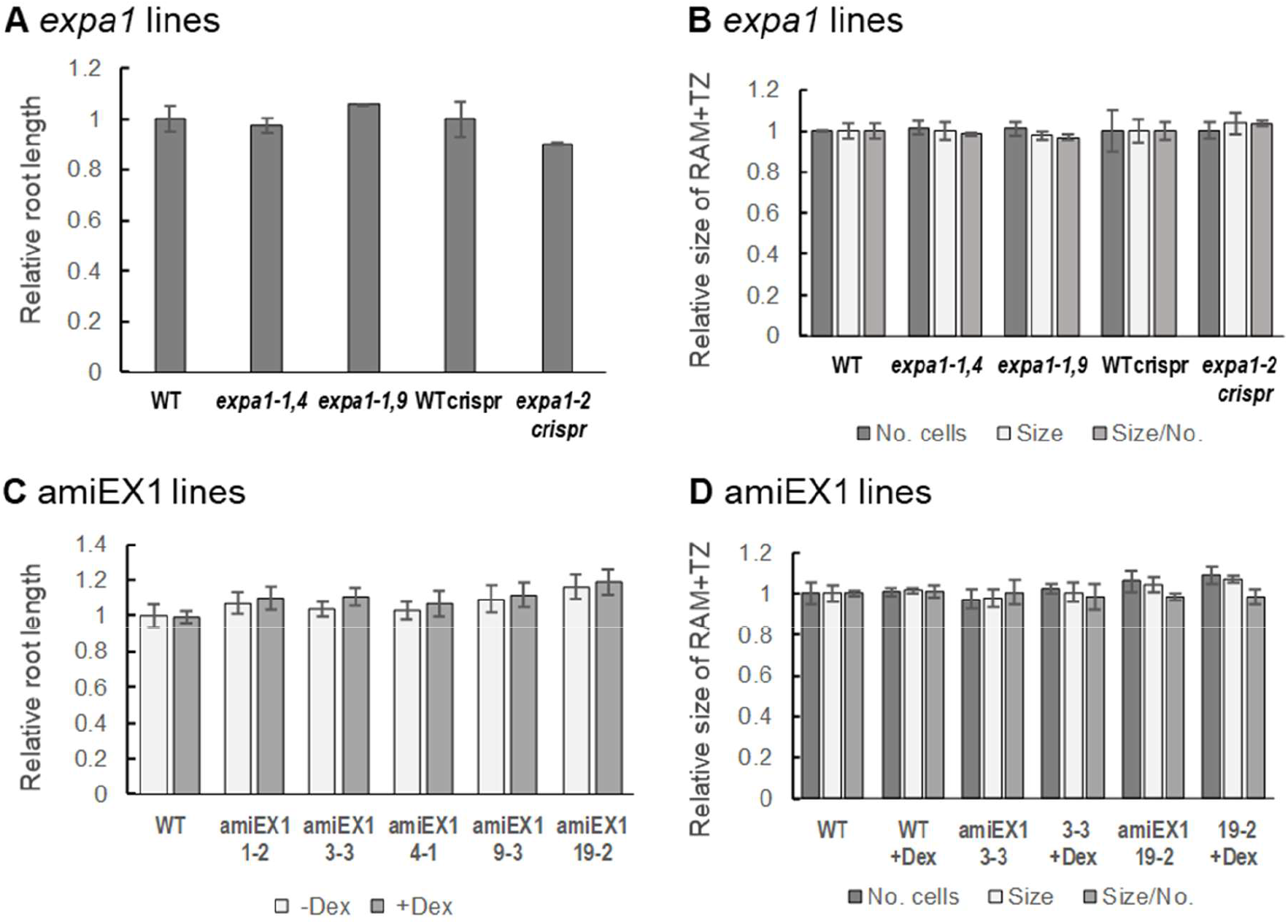
Root growth is not affected in *expa1* knock-out and *EXPA* knock-down lines. **(A)** Root length and **(B)** a number of cells in RAM + TZ, size of RAM + TZ size and the ratio size/ number of cells was measured in 7-day old *Arabidopsis* WT, two families of homozygous knockout lines *expa1-1,4* and *expa1-1,9* (Pacifici *et al*., 2018), CRISPR/Cas9 *expa1-2* (Ramakrishna *et al*., 2019) and its null segregant WT control (WT crispr). **(C)** Root length and **(D)** a number of cells in RAM + TZ, size of RAM + TZ and the ratio size/ number of cells was measured in WT and selected independent T3 homozygous p35S>GR>amiRNA EXPA1, 14, 15 (amiEX1) lines (1-2, 3-3, 4-1, 9-3, 19-3) grown on MS media +/- Dex for 7 days. The data are normalised to the corresponding WT, or WT with mock DMSO treatment (-Dex). The experiment was repeated twice with minimum of 10 seedlings in each category, error bars represent standard error of mean. There are no statistically significant differences.

## SUPPLEMENTARY MATERIALS AND METHODS

### Plasmolysis experiment

*Arabidopsis* seedlings of a plasma membrane marker PM-YFP pUBQ10::YFP-PIP1;4 (von Wangenheim *et al*., 2016) were immersed into 10% solution of sorbitol for cca 10 min.

### EXPA1 mutant lines

Knock-out plants of *EXPA1* were obtained from the Nottingham Arabidopsis Stock Centre collection (SALK_010506). Homozygous mutant lines from the Salk T-DNA were identified by PCR as described (http://signal.salk.edu/) in the next generation of seedlings and designated as *exp1-1,4* and *exp1-1,9*. A second mutant line *exp1-2* generated using the CRISPR/Cas9 and its corresponding WT (WT crispr) was a gift from Alexis Maizel (Ramakrishna *et al*., 2019).

AmiEX1 lines based on artificial microRNAs (amiRNAs, miR319a) were designed using the WMD3 Web MicroRNA Designer (WeigelWorld.org) and the PHANTOM database of family targeting amiRNAs (Hauser *et al*., 2013). The amiRNA sequence engineered for *EXPA1* (*At1g69530*) as well as expansins *EXPA14* (*At5g56320*) and *EXPA15* (*At2g03090*) is “TGTTACACCAACCTGCGGCGT”. We used primers I-IV summarised in Supp. Table 1 to generate PCR fragments that were joined together by overlapping PCR using pRS300 vector as a template (see WeigelWorld.org for details). The final product was cloned into the Dex-inducible pOp6/LhGR vector pOpOn2.1 (Craft *et al*., 2005; Samalova *et al*., 2019) using primers V and VI and GATAWAY™ cloning strategy. Seedlings of selected T3 homozygous lines were used in the experiments.

**Supp. Table 1:**
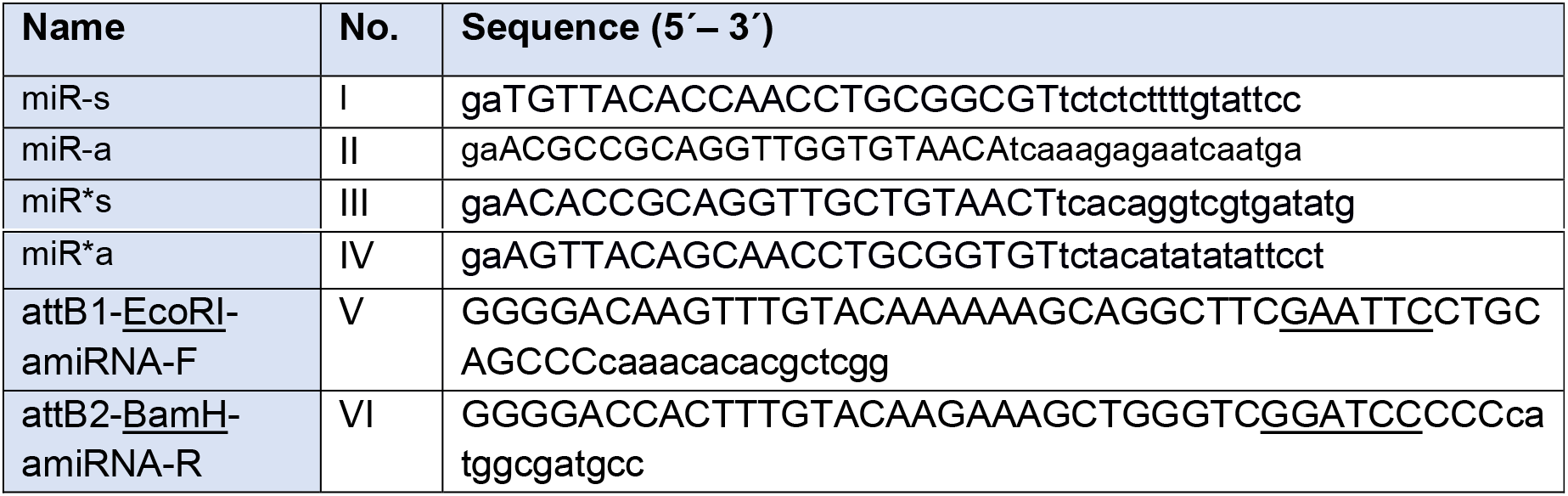
Primes used for cloning of amiEX1 (p35S>GR>amiRNA EXPA1, 14, 15)

